# A Helicase-tethered ORC Flip Enables Bidirectional Helicase Loading

**DOI:** 10.1101/2021.10.06.463322

**Authors:** Shalini Gupta, Larry J. Friedman, Jeff Gelles, Stephen P. Bell

## Abstract

Replication origins are licensed by loading two Mcm2-7 helicases around DNA in a head-to-head conformation poised to initiate bidirectional replication. This process requires ORC, Cdc6, and Cdt1. Although different Cdc6 and Cdt1 molecules load each helicase, whether two ORC proteins are required is unclear. Using colocalization single-molecule spectroscopy combined with FRET, we investigated interactions between ORC and Mcm2-7 during helicase loading. We demonstrate that a single ORC molecule can recruit both Mcm2-7/Cdt1 complexes via similar interactions that end upon Cdt1 release. Between the first and second helicase recruitment, we observe a rapid change in interactions between ORC and the first Mcm2-7. In quick succession ORC breaks the interactions mediating first Mcm2-7 recruitment, releases from its initial DNA-binding site, and forms a new interaction with the opposite face of the first Mcm2-7. This rearrangement requires release of the first Cdt1 and tethers ORC as it flips over the first Mcm2-7 to form an inverted Mcm2-7-ORC-DNA complex required for second-helicase recruitment. To ensure correct licensing, this complex is maintained until head-to-head interactions between the two helicases are formed. Our findings reconcile previous observations and reveal a highly-coordinated series of events through which a single ORC molecule can load two oppositely-oriented helicases.

## Introduction

The eukaryotic DNA replication machinery first assembles and initiates synthesis at DNA sites called origins of replication. During G1, all potential origins are licensed by loading two Mcm2-7 replicative DNA helicases around the origin DNA in an inactive, head-to-head fashion (Abid Ali et al., 2017; Evrin et al., 2009; Li et al., 2015; Remus et al., 2009). This conformation prepares the helicases to initiate bidirectional replication upon activation during S-phase. The Mcm2-7 complex is the core enzyme of the eukaryotic replicative helicase and its loading onto DNA is restricted to G1 phase (Aparicio et al., 1997; Diffley et al., 1994). This constraint prevents helicases from being loaded onto replicated DNA, ensuring that no part of the genome is replicated more than once per cell cycle (Siddiqui et al., 2013).

Interactions between three proteins and the Mcm2-7 helicase direct eukaryotic helicase loading (Bell and Labib, 2016). The origin-recognition complex (ORC) binds origin DNA and then recruits Cdc6 (Bell and Stillman, 1992; Speck et al., 2005). The resulting ORC-Cdc6 complex encircles the origin DNA (Feng et al., 2021; Schmidt and Bleichert, 2020). Mcm2-7 in complex with Cdt1 associates with ORC/Cdc6 and adjacent DNA to form the short-lived <u>O</u>RC-<u>C</u>dc6-<u>C</u>dt1-Mcm2-7 (OCCM) complex (Sun et al., 2013; Ticau et al., 2015). Loading of a second Cdt1-bound-Mcm2-7 hexamer, oriented in the opposite direction of the first, completes formation of the Mcm2-7 “double hexamer”.

Multiple mechanisms have been proposed to explain how two oppositely-oriented helicases are loaded at an origin. One mechanism proposes that distinct ORC molecules bound at inverted DNA sites recruit the two helicases. This model is supported by evidence that mutations in the Mcm3 C-terminus predicted to interfere with interactions between Mcm2-7 and the ORC-Cdc6 complex prevent recruitment of both the first and second helicase (Coster and Diffley, 2017; Frigola et al., 2013). In addition, two ORC DNA binding sites are required for successful helicase loading *in vitro* and origin function *in vivo.* In contrast, single-molecule helicase loading experiments showed that a single ORC molecule can direct loading of both Mcm2-7 helicases (Ticau et al., 2015). A single-ORC model is also supported by time-resolved EM experiments showing predominantly one ORC molecule bound to the DNA in each helicase-loading intermediate observed (Miller et al., 2019). A goal of the current studies is to address these apparently contradictory observations.

Structural studies have revealed important intermediates in helicase loading. ORC has been shown to bend DNA prior to Mcm2-7 recruitment (Li et al., 2018). Cryo-EM studies of the OCCM intermediate show that the first Mcm2-7 hexamer interacts with and encircles the DNA adjacent to ORC-Cdc6 (Sun et al., 2013; Yuan et al., 2017). These and related structures also reveal that interaction of the C-terminal region of ORC with the C-terminal region of Mcm2-7 mediates recruitment of the first Mcm2-7 hexamer (Yuan et al., 2020). ATP hydrolysis is required to proceed beyond the OCCM, and structures of intermediates on-pathway to the Mcm2-7 double hexamer had been elusive until a recent time-resolved cryo-EM study (Miller et al., 2019). Intriguingly, this study observed a complex in which an inverted ORC is engaged with the N-terminal region of the first Mcm2-7. Although it was suggested that formation of the inverted complex might be required to load double hexamers, whether one or two ORC molecules were required to form the complex and load the two Mcm2-7 helicases remained unresolved. Further, this study could not directly examine how the inverted complex was integrated into the sequence of events in helicase loading. A second goal of the current studies is to ask if a single ORC protein can mediate formation of all these stable intermediates and to understand how transitions between the intermediates are coordinated during helicase loading.

Single-molecule biochemical studies have provided kinetic and mechanistic insights into helicase loading that complement structural studies. Single-molecule studies demonstrated that the two Mcm2-7 complexes in each double-hexamer associate with origin DNA in a one-at-a-time manner (Ticau et al., 2015). Single-molecule FRET (sm-FRET) has been used to monitor opening and closing of the interface between Mcm2-Mcm5 (Bochman and Schwacha, 2008; Samel et al., 2014) that provides DNA access to the central channel of the Mcm2-7 ring (Ticau et al., 2017). These studies revealed that Mcm2-7 is recruited with an open gate that closes around origin DNA substantially after initial DNA association. Such approaches also revealed that recruitment and loading of each Mcm2-7 involves a separate set of Cdc6 and Cdt1 molecules. Following each Mcm2-7 recruitment, Cdc6 and Cdt1 are released sequentially, and the Mcm2-7 ring closes concomitant with Cdt1 release (Ticau et al., 2015, 2017). This connection between Cdt1 release and ring closing is consistent with structural data that suggest that Cdt1 holds the Mcm2-7 ring open at the Mcm2-5 gate (Frigola et al., 2017; Zhai et al., 2017).

The mechanism that drives recruitment of head-to-head Mcm2-7 hexamers remains unclear (Bell and Labib, 2016; Lewis and Costa, 2020). Here we generate data supporting a model that reconciles the apparently inconsistent observations regarding ORC and helicase loading and directly observe how a single ORC guides double-hexamer formation. We monitor ORC-Mcm2-7 interactions in real-time using sm-FRET and show that recruitment of each Mcm2-7 hexamer is accompanied by a short ‘OM’ interaction with the same ORC protein. Prior to recruiting the second Mcm2-7 hexamer, ORC forms a distinct intermediate (referred to as ‘MO’) with the initially loaded Mcm2-7. The transition between the OM and MO intermediates is rapid and requires Cdt1 release. Forming the MO intermediate allows ORC to release from its initial binding site and flip over the initially loaded Mcm2-7, positioning ORC to rebind DNA at an inverted binding site without release into solution. The resulting MO intermediate recruits the second Mcm2-7 and is only disrupted when Mcm2-7 double-hexamer formation is initiated. Our findings reveal a highly coordinated series of events that ensures two Mcm2-7 helicases are loaded as head-to-head pairs poised to initiate bidirectional replication.

## Results

### Monitoring ORC-Mcm2-7 interactions during helicase recruitment

To investigate the dynamics of ORC-Mcm2-7 interactions during helicase loading, we developed a sm-FRET assay for the initial interaction between these proteins based on previously described single-molecule helicase-loading experiments (Ticau et al., 2015). Using the structure of the OCCM as a guide (Yuan et al., 2017), we modified ORC and Mcm2-7 at sites that are proximal (∼35 Å apart) during recruitment of the first Mcm2-7 (Figure 1a). ORC was labeled with a donor fluorophore at the Orc5 C-terminus (ORC^5C-549^) and Mcm2-7 was labeled with an acceptor fluorophore at the Mcm2 C-terminus (Mcm2-7^2C-649^). Importantly, the fluorescent labels did not interfere with protein function in ensemble helicase-loading assays (Figure 1-figure supplement 1). To monitor ORC-Mcm2-7 interactions during loading, purified ORC^5C-549^, Mcm2-7^2C-649^, Cdt1, and Cdc6 were incubated with fluorescently-labeled surface-tethered origin-DNA. Using total-internal-reflection fluorescence (TIRF) microscopy, we monitored the colocalization of the fluorescently-modified proteins with individual DNA molecules (Friedman and Gelles, 2015). Alternate excitation of the donor and acceptor fluorophores (Figure 1-figure supplement 2b) allowed observation of the association of each labeled protein with origin DNA, and determination of the apparent FRET efficiency (*E*_FRET_) during donor excitation measured ORC-Mcm2-7 C-terminal interactions (Figure 1a and 1b). We will refer to the ORC-Mcm2-7 interactions monitored using FRET between ORC^5C-549^ and Mcm2-7^2C-649^ as “OM interactions.”

**Figure 1.**
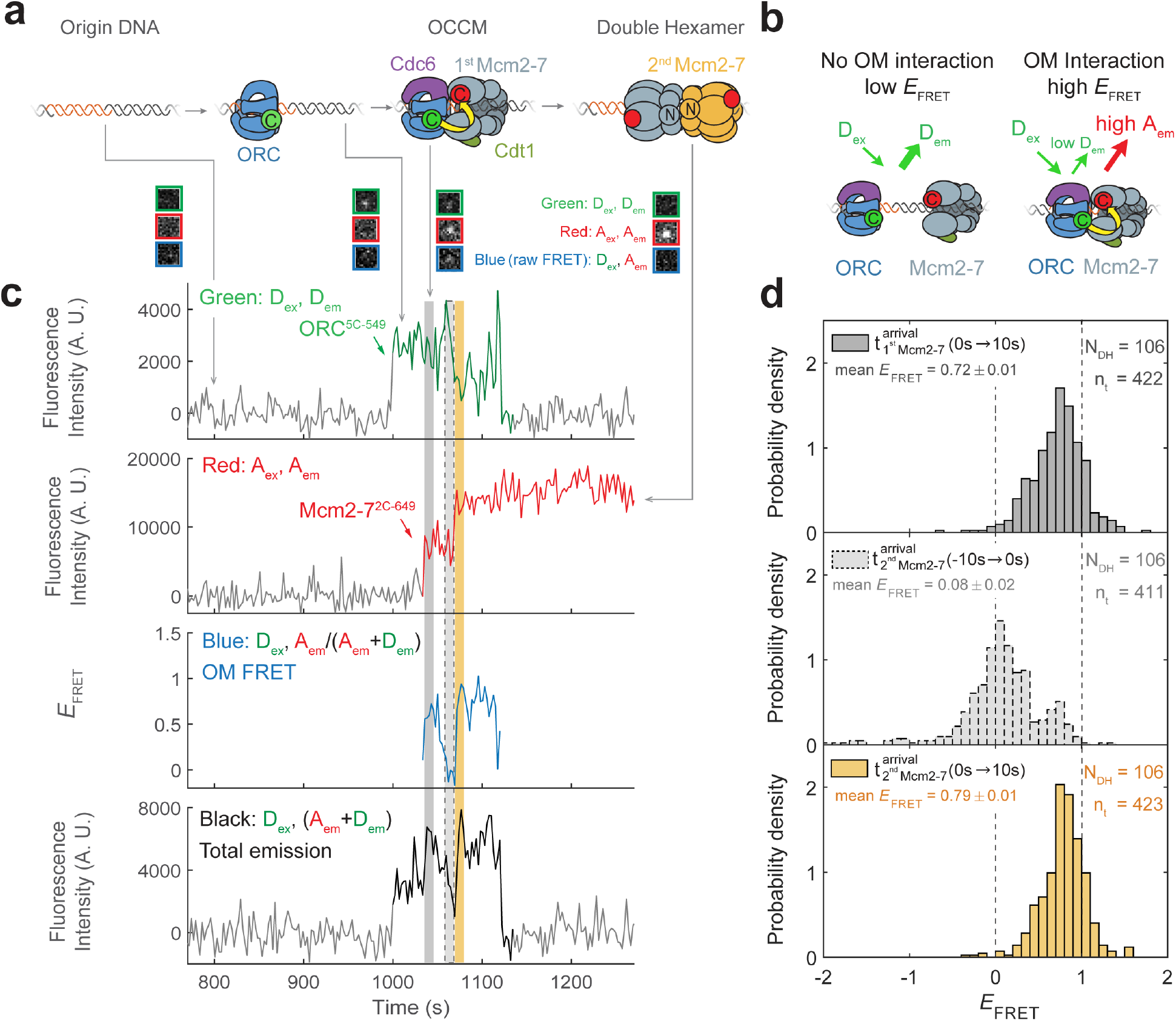
ORC recruits two Mcm2-7 helicases through sequential OM interactions. **(a)** A schematic of the molecular events during helicase loading. The ORC and Mcm2-7 proteins are labeled at their C-termini (C) which form an interface during first Mcm2-7 recruitment (Yuan et al., 2017). ORC^5C-549^ is labeled with a donor fluorophore (D; green circles) and Mcm2-7^2C-649^ is labeled with an acceptor fluorophore (A; red circles). The associated fluorescent images show single frames of ORC, Mcm2-7 and raw FRET fluorescence spots at a single DNA molecule in green, red, and blue outlines, respectively. We use gray and yellow coloring to indicate the first and second Mcm2-7 hexamers respectively. **(b)** Schematic of DNA-bound ORC^5C-549^ and Mcm2-7^2C-649^ proteins. When ORC^5C-549^ and Mcm2-7^2C-649^ are not associated, excitation of the donor (D_ex_) only yields emission from the donor fluorophore (D_em_). However, when ORC^5C-549^ and Mcm2-7^2C-649^ are in proximity, we observe emission from the acceptor (A_em_) on donor excitation due to FRET, and corresponding lower emission from the donor fluorophore (D_em_). **(c)** A representative trace showing ORC and Mcm2-7 associations with DNA and ORC-Mcm2-7 (OM) interactions during helicase loading. Donor-excited fluorescence record shows ORC^5C-549^ association (Green: D_ex_, D_em_ panel) and acceptor-excited fluorescence record shows Mcm2-7^2C-649^ association (Red: A_ex_, A_em_ panel). Gray arrows link to the corresponding molecular events shown in (a). Interaction between ORC and Mcm2-7 decreases the distance between the donor and acceptor fluorophores and results in increased FRET efficiency (*E*_FRET_, Blue: D_ex_, A_em_/ (A_em_ + D_em_) panel). The black: D_ex_, (A_em_ + D_em_) panel shows donor-excited total emission (see Figure 1–figure supplement 3). An objective image-analysis algorithm (Friedman and Gelles, 2015) detects a spot of fluorescence at time points shown in green, red, and black on the time records. *E*_FRET_ values are shown for time points at which both D and A are present. Gray, gray-dash and yellow highlight three 10 second (s) time intervals referenced in (d). A.U., arbitrary units. Concentrations of labeled proteins in the reaction are 0.5 nM ORC^5C-549^, 15 nM Cdt1-Mcm2-7^2C-649^. Figure 1–figure supplement 2a shows additional records. **(d)** Histogram plots of *E*_FRET_ values for 106 single-ORC-mediated, double-hexamer formation events during three 10 s time intervals: immediately after the first (top) or second Mcm2-7 (bottom) arrives or before the second Mcm2-7 arrives (middle). Examples of these intervals are indicated in (c) with gray, yellow, and gray dash, respectively. The two dashed lines indicate *E*_FRET_ values of 0 and 1. N_DH,_ number of double-hexamer formation events and n_t_, number of signal points. Rare *E*_FRET_ values below -2 and above +2 were excluded (2/428, 13/428 and 1/428 signal points from the top, middle and bottom histograms respectively).

We focused our studies on SM event records that are consistent with Mcm2-7 double hexamer formation. Mcm2-7 complexes associated with DNA in a one-at-a-time fashion as described previously (Ticau et al., 2015). A fraction of the DNAs (∼28% of total DNAs) showed two stepwise increases in Mcm2-7^2C-649^ fluorescence intensity as expected for formation of the Mcm2-7 double hexamer. Of the DNAs with two Mcm2-7 associations, we limited further analysis to a subset showing two Mcm2-7 complexes that are retained on DNA for 20 or more frames of acquisition (>48 s). These long-lived sequential associations occurred on 12% of DNAs and represent successful Mcm2-7 double-hexamer formation (Ticau et al., 2015).

Monitoring ORC and Mcm2-7 DNA association showed that for most double-hexamer formation events, one ORC molecule loaded two Mcm2-7 complexes. Specifically, we observed association of one ORC molecule with the DNA throughout the interval for sequential recruitment of two Mcm2-7 molecules in 81% of double-hexamer formation events. The observation of only one ORC is not due to incomplete ORC labeling as we determined that 88% ± 2% of the ORC^5C-549^ protein was labeled (see Methods). Thus, if a second ORC was required for recruiting the two helicases, we would expect to see association of two ORCs in 77% (0.88^2^ = 0.77) of double-hexamer formation events. In contrast, we observed two ORC proteins during sequential Mcm2-7 recruitment in only 13% of the double-hexamer formation events (22/166). The remaining 6% of double-hexamer formation events had three associated ORC molecules, likely reflecting one or more non-specific ORC DNA binding events.

In the small fraction of double-hexamer formation events with two ORC molecules, the arrival of the second ORC was not coordinated with any other event we could observe during helicase loading. We found three types of two-ORC double-hexamer formation events. In 8/166 events, separate ORC molecules were present during the two Mcm2-7 associations. The first ORC appeared to release from DNA following the recruitment of the first Mcm2-7, and a second ORC was present during the recruitment of the second Mcm2-7. These rare events provide the first direct evidence of helicase loading mediated by two separate ORC molecules. For 14/166 events, a second ORC associated between the first and second Mcm2-7 recruitment events. We note that for this category, it is possible that a single ORC molecule mediates both recruitment events with the second ORC being DNA-bound but uninvolved (see ‘The same ORC molecule interacts with both Mcm2-7 helicases’). We restricted subsequent analysis to events with a single ORC associated throughout both helicase recruitment events for two reasons: they were the most frequent events observed (81% of the successful helicase loading events), and to investigate how one ORC could load two Mcm2-7 helicases in opposite orientations.

FRET between ORC^5C-549^ and Mcm2-7^2C-649^ is a reporter for C-terminal ORC-Mcm2-7 interactions during helicase recruitment (Figure 1b). When both ORC^5C-549^ and Mcm2-7^2C-649^ were co-localized with DNA, we observed transient periods of high *E*_FRET_ upon Mcm2-7 DNA association (Figure 1c, Blue: D_ex_, A_em_/(A_em+_D_em_) panel). Mean *E*_FRET_ ± S.E.M. in the 10 s intervals after first and second Mcm2-7 arrivals were 0.72 ± 0.01 and 0.79 ± 0.01 respectively (Figure 1d, top and bottom panels). Three findings give us confidence that these high *E*_FRET_ values arise from interactions between ORC and Mcm2-7. First, we positioned the fluorescent probes on ORC and Mcm2-7 to detect interactions between regions that are proximal in the complex presumed to represent the initially recruited Mcm2-7, the OCCM (Yuan et al., 2017). Second, in 105/111 instances where a first Mcm2-7^2C-650^ arrived at an ORC^5C-549^-bound-DNA, high *E_FRET_* was detected at the same time as the arrival of the Mcm2-7 on DNA (temporal resolution is ±2.4 s; Figure 1– figure supplement 2b and 2c). This coordination indicates that the high *E*_FRET_ value monitors an interaction occurring during initial Mcm2-7 recruitment. Finally, we verified that observing high *E*_FRET_ was dependent on the placement of the fluorescent probes on Mcm2-7 and ORC. Moving the label on either ORC or Mcm2-7 to the opposite side of the protein relative to the C-terminal ORC-Mcm2-7 interface seen in the OCCM resulted in strong reductions in the associated *E*_FRET_ values (Figure 2–figure supplement 1, Figure 3–figure supplement 1). Together, these observations establish that the high *E*_FRET_ state arises from specific ORC-Mcm2-7 interactions during initial recruitment of Mcm2-7.

**Figure 2.**
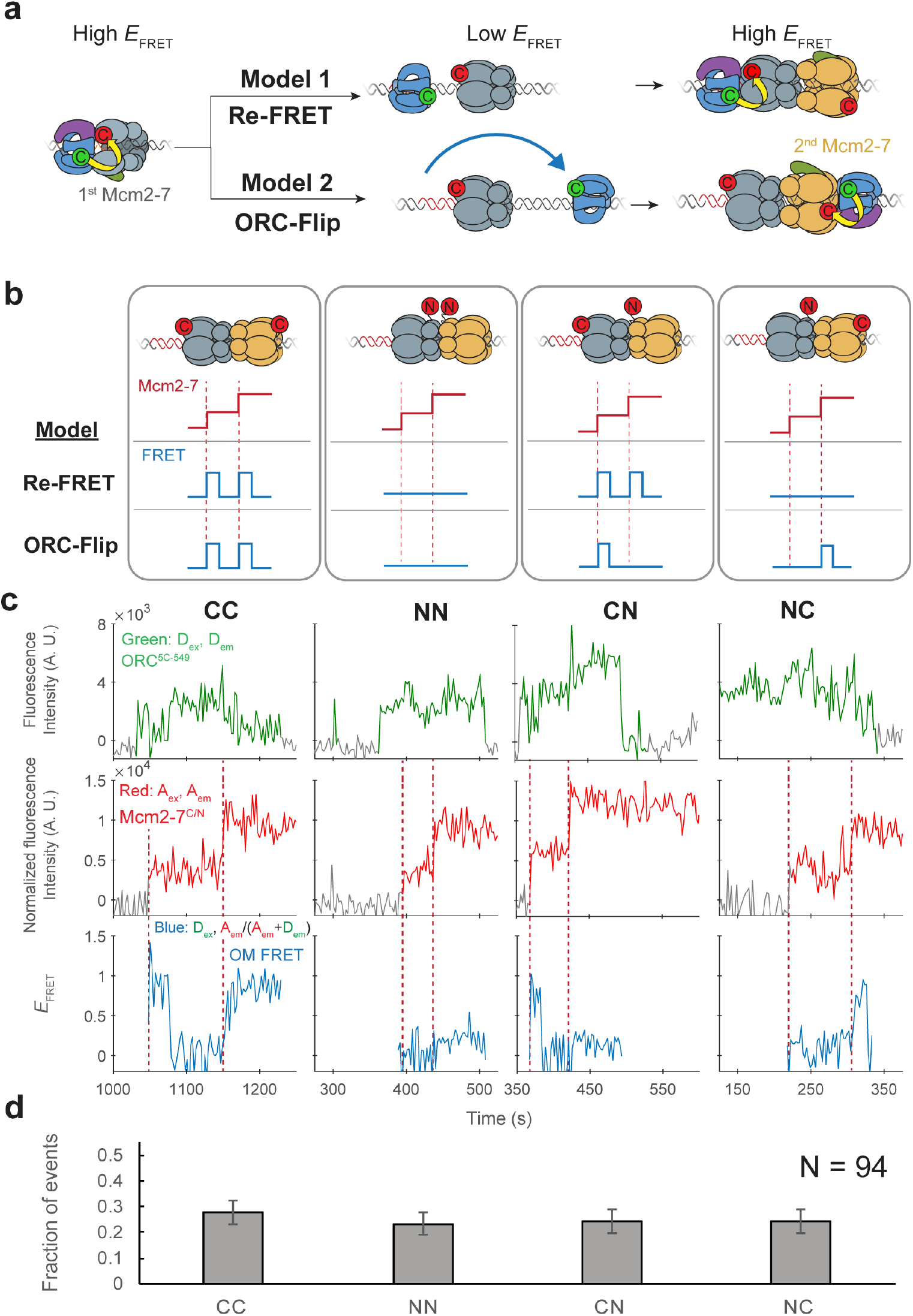
ORC makes OM interactions sequentially with the first and second Mcm2-7 helicases. **(a)** Two models to explain an OM-FRET interaction accompanying each Mcm2-7 arrival in an experiment (Figure 1) in which both ORC and Mcm2-7 are C-terminally labeled. Both models begin with ORC recruiting the first Mcm2-7 during the first OM interaction (high *E*_FRET_). In the Re-FRET model, ORC separates from the first Mcm2-7 (resulting in low *E*_FRET_) followed by re-interacting with the first Mcm2-7. In the ORC-flip model, ORC releases from its original binding site and rebinds the DNA at a second inverted binding site on the other side of Mcm2-7, resulting in lower *E*_FRET_. The flipped ORC then recruits the second Mcm2-7 via a second OM interaction. **(b)** Experimental setup to distinguish between the Re-FRET and ORC-Flip models. Acceptor-labeled Mcm2-7^2C-649^ (C; red circles) and Mcm2-7^4N-650^ (N; red circles) are mixed in an equimolar ratio, resulting in four subpopulations of double hexamers – CC, NN, CN and NC. Red dashed lines indicate Mcm2-7 arrival times during double-hexamer formation. Only the C-terminally labeled Mcm2-7^2C-649^ molecules exhibit high FRET with donor-labeled ORC^5C-549^. The Re-FRET and ORC-flip models predict distinct FRET profiles for the four double hexamer populations generated. Importantly, the observation of single FRET peaks is unique to the ORC-flip model. **(c)** Representative double-hexamer formation events from an experiment with mixed Mcm2-7^2C-649^ and Mcm2-7^4N-650^. Concentrations of labeled proteins in the reaction are 0.5 nM ORC^5C-549^, 7.5 nM Cdt1-Mcm2-7^2C-649^ and 7.5 nM Cdt1-Mcm2-7^4N-650^. Figure 2–figure supplement 2 shows additional records of CN and NC double hexamers. **(d)** The fraction of observed events (± standard error, SE) corresponding to the type of FRET profile shown above in (c).

**Figure 3.**
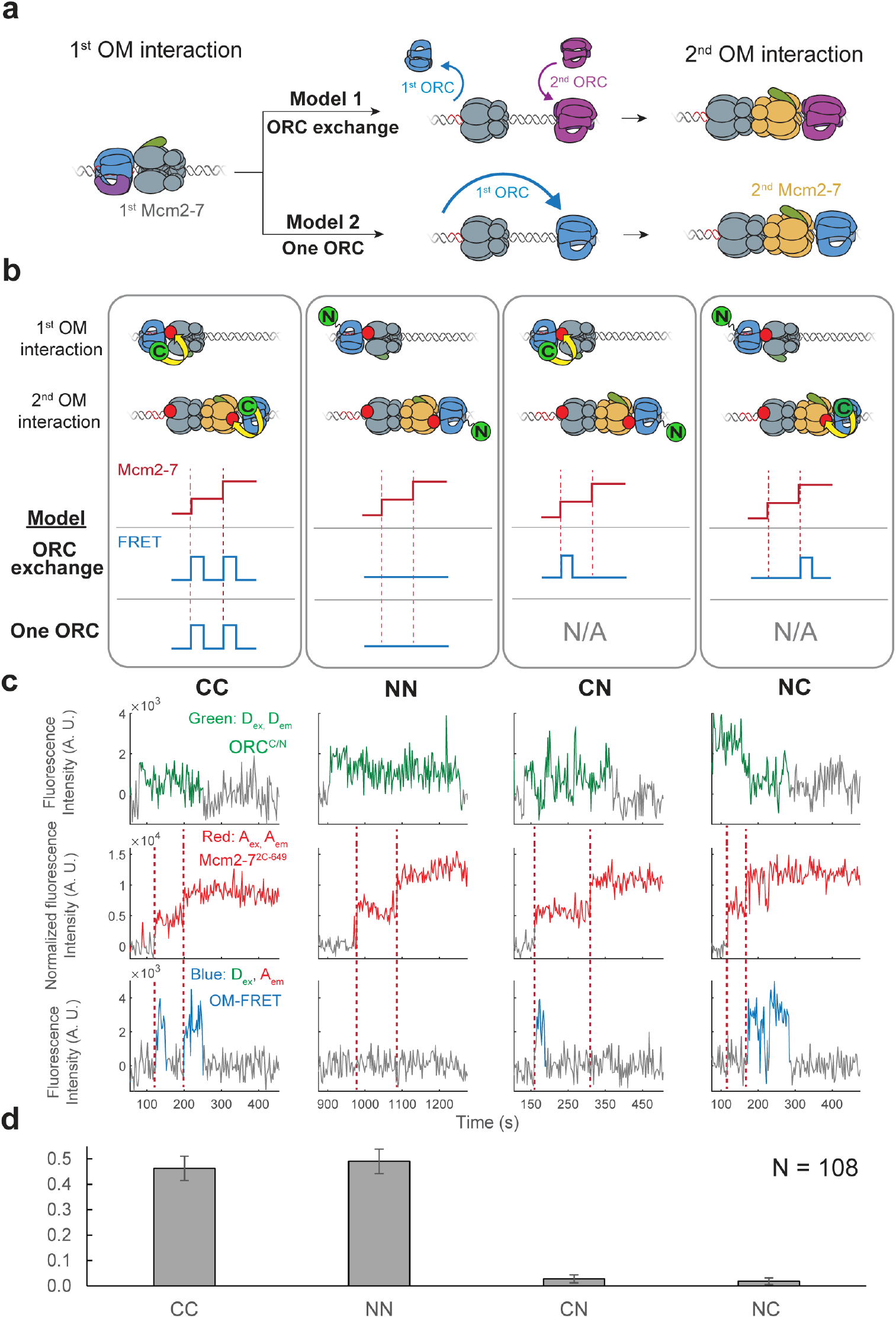
The same ORC molecule interacts with both Mcm2-7 helicases. **(a)** Two models that can explain how ORC interacts with both Mcm2-7 helicases. Both models are consistent with the observation of uninterrupted fluorescence of a single ORC throughout both helicase recruitment events. In the ‘ORC-exchange’ model, what appears to be a single, continuously present ORC is in fact rapid and coordinated exchange of two ORC molecules that recruit the first and second Mcm2-7 helicases, respectively. In the ‘one-ORC’ model, the same ORC recruits both helicases. **(b)** Experimental setup to distinguish between the ORC-exchange and one-ORC models. A mixture of donor-labeled ORC^5C-549^ (C; green circles) and ORC^1N-550^ (N; green circles) is added to a reaction with acceptor labeled Mcm2-7^2C-649^. Only ORC^5C-549^ (C-terminally labeled) can FRET with Mcm2-7^2C-649^ (Figure 3–figure supplement 1). Red dashed lines indicate Mcm2-7 arrival times. The predicted FRET results are illustrated for each model. There are four possible ORC pairs and FRET results when two ORC molecules mediate first and second OM interaction – CC, NN, CN and NC. However, in the one-ORC model only the CC and NN pairs are possible resulting in only two possible FRET results. N/A, not applicable. **(c)** Representative double hexamer formation events in an experiment with mixed ORC^5C-549^ and ORC^1N-550^. Concentrations of labeled proteins in the reaction are 0.25 nM ORC^5C-549^, 0.60 nM ORC^1N-550^ (see methods) and 15 nM Cdt1-Mcm2-7^2C-649^. **(d)** The fraction of events that correspond to each type of FRET profile shown above in (c).

We note that the total emission upon donor excitation is enhanced during OM interactions (Figure 1c, Black: Dex, Dem + Rem panel). This is due to protein-induced fluorescence enhancement (Hwang et al., 2011) of the of the DY549P1 fluorophore during the OM interaction (Figure 1–figure supplement 3). Importantly, this effect does not hinder our ability to detect OM interactions. Instead, it represents an additional indicator of this interaction.

### Each Mcm2-7 recruitment is accompanied by an OM interaction

Monitoring the ORC-Mcm2-7 recruitment interface revealed two periods of OM interaction during helicase loading. Arrival of each of the two Mcm2-7 helicases was accompanied by a short period of high *E*_FRET_ in almost all cases (Figure 1c, Blue: D_ex_, A_em_/(A_em+_D_em_) panel). In 111 examples of double-hexamer formation, 106 showed distinct periods of high *E*_FRET_ with each of the two Mcm2-7 arrivals (see below for a description of the remaining 5/111 cases). The elevated *E*_FRET_ values were simultaneous with the arrival of the corresponding Mcm2-7 on the DNA (Figure 1–figure supplement 2c), consistent with both OM interactions mediating Mcm2-7 recruitment. The start time of the first OM interaction matched the first Mcm2-7 arrival in 104/106 cases and the start of the second OM interaction matched the arrival of the second Mcm2-7 in 98/106 cases (time resolution of experiment 2.4 s). The rare cases (2/106 and 7/106 for the first and second Mcm2-7 respectively) that were not simultaneous were all within 4.8 s of the corresponding Mcm2-7 arrival except for one (6 s). The similar mean *E*_FRET_ values associated with the first and second OM interactions (0.72 ± 0.01 and 0.79 ± 0.01, respectively; Figure 1d) indicate that recruitment of the first and second Mcm2-7 is mediated by the formation of a similar ORC-Mcm2-7 interface.

The first OM interaction consistently ended before the second Mcm2-7 was recruited during helicase loading. Although both ORC and first Mcm2-7 remained associated with the DNA, the OM-FRET signal was lost before second Mcm2-7 arrival in 106/106 cases. Thus, there was always a period lacking OM interaction between the two Mcm2-7 recruitment events. Consistent with this observation, *E*_FRET_ values in the 10 s interval preceding the second Mcm2-7 arrival were low (Figure 1d, middle panel), with an average value of 0.08 ± 0.02. The second OM interaction observed was also short-lived and in a majority of events ends with the loss of ORC^5C-549^ fluorescence (71/106; Figure 1–figure supplement 4a). We confirmed this loss of donor fluorescence was typically not due to photobleaching (Figure 1–figure supplement 4b), indicating that the end of the second OM interaction in these cases was concomitant with the departure of ORC from the DNA. These data indicate that the interactions that recruit the first Mcm2-7 are always broken before the second Mcm2-7 is recruited, as expected if the same ORC mediated both events.

### ORC interacts with both the first and the second Mcm2-7 during double-hexamer formation

Two distinct mechanisms of helicase loading could result in the observation of two sequential OM interactions with a single ORC protein. The two high *E*_FRET_ signals could arise due to ORC^5C-549^ interacting twice with the first Mcm2-7^2C-649^ (Figure 2a Model 1, ‘Re-FRET’). Alternatively, ORC^5C-549^ may instead interact with each of the two Mcm2-7^2C-649^ complexes on arrival at the DNA (Model 2). To achieve the head-to-head conformation of the Mcm2-7 helicases in the double-hexamer, Model 2 would require either ORC or the first Mcm2-7 to invert its orientation on the DNA. Since initial loading of the Mcm2-7 complex involves passing origin DNA through the Mcm2-5 gate followed by gate closing, inversion of Mcm2-7 on the DNA would require re-opening of the Mcm2-5 gate. However, previous studies found no evidence of Mcm2-5 gate re-opening during helicase loading (Ticau et al., 2017). For this reason, the version of Model 2 in which ORC rebinds the DNA on the opposite side of the first Mcm2-7 and in the opposite orientation is most likely. We call this the ‘ORC-flip’ model.

To distinguish between the Re-FRET and ORC-flip models, we performed the SM helicase-loading reaction with a mixture of two Mcm2-7 preparations labeled at different positions with acceptor fluorophores (Mcm2-7^2C-649^ and Mcm2-7^4N-650^). Importantly, in contrast to Mcm2-7^2C-649^, the N-terminally labeled Mcm2-7^4N-650^ exhibits very low FRET with ORC^5C-549^ (Figure 2–figure supplement 1). Mixing an equimolar ratio of these two modified Mcm2-7 complexes with ORC^5C-549^ allows the formation of four distinct populations of double-hexamers (Figure 2b). We will refer to the four populations of double hexamers by the position of the acceptor fluorophore on the first and second helicase in that order. Thus, a double hexamer formed where the first helicase is Mcm2-7^2<u>C</u>-649^ and second is Mcm2-7^4<u>N</u>-650^ will be referred to as ‘CN’ and a double hexamer where the first helicase is Mcm2-7^4<u>N</u>-650^ and second is Mcm2-7^2<u>C</u>-649^ is ‘NC’. Double hexamers with the same labeled Mcm2-7 for the first and second events would be ‘CC’ or ‘NN.’

Depending on which mechanism is used, the four double-hexamer populations will generate distinct OM interaction profiles (Figure 2b). If the Re-FRET model is accurate, ORC would only interact with the first recruited helicase. Since the CC and CN double hexamers have Mcm2-7^2C-649^ as the first helicase, these populations should have two sequential high FRET peaks corresponding to each helicase arrival. In contrast, because the NN and NC double hexamers have Mcm2-7^4N-650^ as the first helicase, the Re-FRET model predicts these populations should not exhibit high FRET during double-hexamer formation.

If the ORC-flip model is correct, ORC would form an OM interaction sequentially with each Mcm2-7 helicase in the double hexamer. Thus, for this model only CC double hexamers should be associated with two sequential high FRET peaks. Importantly, for the ORC-flip model the mixed double hexamers (CN and NC) would exhibit a single high FRET peak when the C-terminally labeled Mcm2-7^2C-649^ arrives. Thus, the observation of single OM interaction *E*_FRET_ (OM-FRET) peaks during loading of two helicases is unique to the ORC-flip model, allowing us to distinguish between the two models.

We observed four distinct patterns of OM interaction profiles at similar frequencies when the differently-labeled Mcm2-7 complexes were present (Figure 2c, Figure 2–figure supplement 2). Of 90 double-hexamer formation events, we identified 25 events with two OM-FRET peaks and 21 events with no associated OM-FRET. Strikingly, ∼half the double hexamer formation events had a single OM-FRET peak associated with them (44/90), consistent with the ORC-flip model (see Figure 2b, CN and NC panels). Also consistent with this model, these single OM-FRET events are equally distributed between events in which FRET is associated with the first (22/90 events) or the second (22/90 events) Mcm2-7 arrival (Figure 2d). Indeed, based on the *E*_FRET_ patterns, we were able to infer the underlying double-hexamer “type” for each OM-FRET pattern revealing similar frequencies of CC, NN, CN, and NC double hexamers. Both the presence of the four patterns of FRET and their equal frequency strongly support the ORC-flip model. We note, that these patterns cannot be explained by incomplete labeling of Mcm2-7 complexes, as we only analyzed events with two labeled Mcm2-7 and a labeled ORC.

In the experiment with only C-terminally labelled Mcm2-7 (Figure 1), most events exhibited two sequential OM-FRET peaks, but rare events did not. We observed 1/111 events in which only the first Mcm2-7 arrival exhibited FRET and 3/111 events in which only the second Mcm2-7 arrival exhibited FRET. We also observed 1/111 events with no FRET on either Mcm2-7 arrival. Although all five events appear to have a single ORC co-localized with DNA, these events are likely to arise from DNAs with two bound ORC molecules, one of which is unlabeled. Importantly, these events occur at very low frequency and cannot explain the frequent observation of one or no FRET peaks observed in reactions with mixed Mcm2-7^2C-649^ and Mcm2-7^4N-650^ complexes (Figure 2d). Thus, we conclude that ORC makes OM interactions with both the first and second Mcm2-7 helicases during helicase loading.

### The same ORC molecule interacts with both Mcm2-7 helicases

Although we saw a single ORC remain co-localized with DNA during helicase loading, it remained possible that a rapid exchange of ORC molecules occurred between the first and second Mcm2-7 recruitment. Two models (Figure 3a) could explain how ORC might interact with both Mcm2-7 helicases on recruitment in a manner that is consistent with our previous results. First, two ORCs could be involved in helicase loading, with a separate ORC molecule guiding the recruitment of each of the two Mcm2-7s. To be consistent with our previous observations, the two ORCs would need to exchange in a rapid and coordinated manner such that only one ORC appears co-localized with DNA at any time. We will refer to this as the ‘ORC-exchange’ model. An alternative model posits that the same ORC interacts with both the first and the second helicase (‘one-ORC’ model). In this case the same ORC would release from its initial DNA binding site and rebind the DNA in the opposite orientation prior to recruitment of the second Mcm2-7.

To distinguish between the ORC-exchange and one-ORC models, we used a mixed-ORC-labeling approach (Figure 3b). Analogous to the mixed-Mcm2-7-labeling used in Figure 2, we performed the helicase-loading reaction with Mcm2-7 labeled in a single position (Mcm2-7^2C-649^) and a mixture of ORC proteins labeled at distinct sites (ORC^5C-549^ and ORC^1N-550^). Only the C-terminally labeled ORC protein shows significant FRET with Mcm2-7^2C-649^ (Figure 3–figure supplement 1). In the ORC-exchange model, four possible pairs of ORC molecules could mediate helicase loading. We will refer to these ORC pairs as CC, NN, CN or NC based on the position of the fluorophore on the first and second ORC molecules, respectively. For example, the mixed ORC pair CN indicates ORC^5<u>C</u>-549^ as the first ORC and ORC^1<u>N</u>-550^ as the second. In contrast, the one-ORC model requires that a single ORC mediate both OM interactions. Thus, for this model, only two of the four possible ORC ‘pairs’ described above (CC and NN) are possible because the same ORC recruits the first and second Mcm2-7.

We observed two predominant patterns of OM interaction profiles in the experiment with the mixed ORC proteins (Figure 3c). Of 108 double-hexamer formation events, 50 events had two OM-FRET peaks (fraction of events is 0.46 ± 0.05; Figure 3d) and 53 events had no associated OM-FRET (0.49 ± 0.05). The profiles with two OM-FRET peaks can only come from CC ORC ‘pairs’ and those with no OM-FRET are characteristic of NN ORC ‘pairs’ (see CC and NN panels in Figure 3b). We identified 3/108 events that were consistent with CN ORC pairs (0.03 ± 0.02) and 2/108 events with NC ORC pairs (0.02 ± 0.01). The ORC exchange model predicts that the fraction of events with CC and NN ORC pairs should be similar to those for the mixed ORC pairs (CN and NC). However, the vast majority of events are of CC and NN type (103/108), which is consistent with the same ORC interacting with both Mcm2-7 helicases. The rare CN and NC events are most likely two-ORC events in which one ORC is unlabeled (see above). Based on these findings, we conclude that the majority of events involve one ORC molecule interacting sequentially with both helicases (one-ORC model).

The above analyses included only events in which one fluorescent ORC was present throughout loading. We also analyzed 27 events in which more than one fluorescent ORC was present during the loading event. In a subset of these events (7/27) the first ORC disappeared before the second one arrived, raising the possibility of a two-ORC mechanism. We also observed 20/27 events in which a second ORC associated between the first and second Mcm2-7 recruitment events while the first ORC remained present throughout. Interestingly, a substantial majority (17/20) of these events showed either the CC or the NN pattern consistent with only the first ORC mediating both recruiting events. These observations suggest that the fraction of helicase loading events mediated by two ORCs is lower than the small fraction in which two ORCs were present.

### Cdt1 release corresponds with the end of OM interaction

Measuring the duration of OM interactions revealed a kinetic difference between the first and second OM interactions. In each case, the OM interactions are relatively short-lived, however, the distributions of first and second events are distinct. A survival plot of the two OM interactions (Figure 4a) shows the distribution of OM interaction time during first Mcm2-7 recruitment is significantly shorter (mean, 30 ± 2 s) than that during recruitment of the second Mcm2-7 helicase (mean, 56 ± 4 s). Interestingly, the average OM interaction durations are similar to the mean dwell times of the first and second Cdt1 molecules reported previously (Ticau et al., 2015). This similarity raised the possibility that the release of Cdt1 that occurs during loading of each Mcm2-7 is connected to the termination of the OM interaction.

**Figure 4.**
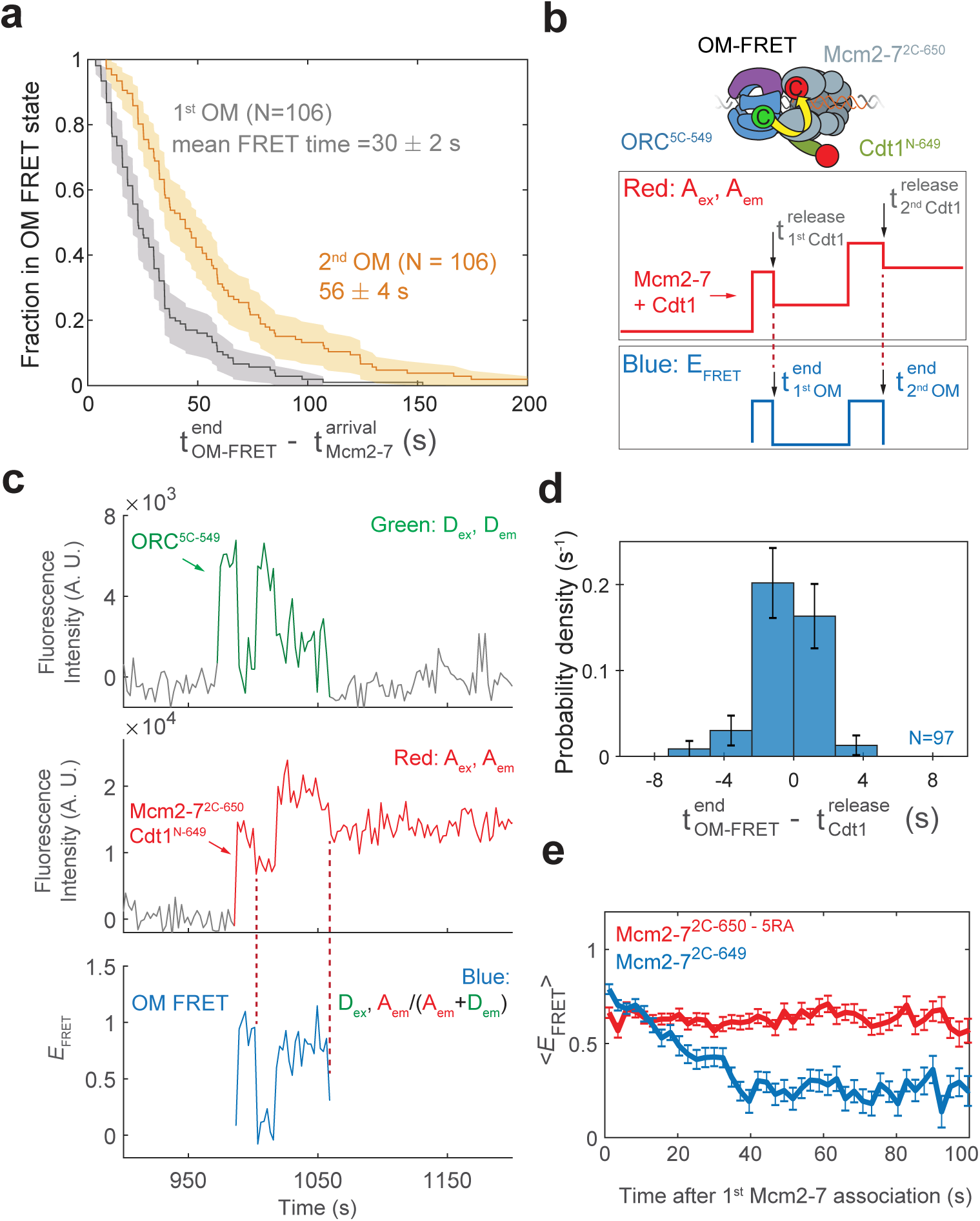
Cdt1 release is concomitant with the end of each OM interaction on individual DNAs. **(a)** Duration of the first (gray) and second (yellow) OM interactions after the association of the corresponding Mcm2-7 (from experiment in Figure 1) are plotted as survival functions. Shaded areas represent the 95% confidence intervals for each curve. **(b)** Experimental setup for observation of Cdt1 release and OM interactions in the same recording. ORC and Mcm2-7 are labeled using the OM-FRET pair at ORC^5C-549^ (donor) and Mcm2-7^2C-650^ (acceptor). Cdt1^N-649^ is also labeled with an acceptor fluorophore, but at a position that exhibits low *E*_FRET_ with ORC^5C-549^ (Figure 4–figure supplement 1). The example traces show the expected results if Cdt1 release is simultaneous with the end of the OM interaction. Each Mcm2-7^2C-650^/Cdt1^N-649^ association results in an increase in fluorescence in the acceptor-excited fluorescence record (Red: A_ex_, A_em_ panel). Black arrows indicate the Cdt1^N-649^ release, which results in loss of roughly half of the acceptor fluorescence. Each Mcm2-7^2C-650^/Cdt1^N-649^ arrival is accompanied by high *E*_FRET_ due to OM interaction (Blue: *E*_FRET_ panel). The black arrows indicate the end of OM-FRET if Cdt1 release is concomitant with the end of OM interactions. **(c)** A representative trace showing the sequential association of two Mcm2-7^2C-650^/Cdt1^N-649^ molecules and OM-FRET. The red dashed lines indicate Cdt1 release times. Concentrations of labeled proteins in the reaction are 0.5 nM ORC^5C-549^, 10 nM Mcm2-7^2C-649^ and 10 nM Cdt1^N-649^. Figure 4–figure supplement 3 shows additional records. **(d)** Cdt1 release is correlated with the end of OM interactions. The duration of OM-FRET after Cdt1 release (±SE) is plotted as a histogram. Data from the first and second Cdt1 releases is combined plotted with a bin size of 2.4 s. We could observe Cdt1 association and release in 99/114 Mcm2-7 association events (Cdt1 was unlabeled in 15/114 events). Not shown are 2/99 events where OM interactions persisted much longer after the corresponding Cdt1 was released (56.8 s and 179.8 s). **(e)** OM interaction times are extended when Cdt1 release is disrupted. Comparison of the population-average OM-interaction *E*_FRET_ values (±SEM) at indicated times after the first Mcm2-7 association for Mcm2-7^2C-650,5RA^ (which is defective in Cdt1 release; N=128) and Mcm2-7^2C-649^ (N=106). In the experiment with Mcm2-7^2C-650,5RA^, sample size was limited to the first half of the DNA locations (174/348), which had 128 Mcm2-7 association events with OM-FRET.

To examine the possibility that Cdt1 release is linked to the end of the corresponding OM interaction, we monitored Cdt1 release and OM-FRET in a single experiment. To this end, in addition to the previously described fluorescently-labeled OM-FRET pair, we included Cdt1 labeled at its N-terminus with a red-excited acceptor fluorophore (Cdt1^N-649^). Mcm2-7 and Cdt1 arrive together at origin DNA (Ticau et al., 2015). Thus, in this experiment each Mcm2-7/Cdt1 arrival results in the association of two red fluorophores with DNA (Figure 4b, Red: A_ex_, A_em_ panel). Successful helicase loading is associated with the release of Cdt1 while Mcm2-7 remains bound to the DNA (Ticau et al., 2015). Such Cdt1 release events will result in the loss of one red fluorophore and appear as the reduction of the acceptor fluorescence intensity (Red: A_ex_, A_em_). Although Cdt1^N-649^ is also labeled with an acceptor fluorophore, this site of labeling does exhibit strong FRET with ORC^5C-549^ (Figure 4–figure supplement 1). In a control experiment in which only Cdt1 is labeled with the acceptor fluorophore, the average *E*_FRET_ of observed when Cdt1 is present was 0.358 ± 0.003, which is easily distinguished from that of the OM-FRET pair (average *E*_FRET_ 0.72 ± 0.01; see Figure 1d). However, the presence of Cdt1^N-649^ does contribute to a slightly elevated *E*_FRET_ when the OM interaction is formed (mean *E*_FRET_ = 0.83 ± 0.01 with Cdt1^N-649^ compared to ∼0.72 with unlabeled Cdt1, Figure 4–figure supplement 1b). Thus, if Cdt1 release is concomitant with OM interactions ending, the loss of half the acceptor fluorescence intensity due to Cdt1 release should occur simultaneously with the loss of FRET (Figure 4b, Blue: *E*_FRET_ panel). If Cdt1 release is disconnected from OM interactions ending, we would observe only a small reduction (∼0.83 to 0.72) in *E*_FRET_ when Cdt1^N-649^ is released.

We observed a clear correlation between the time that OM interactions end and the time of Cdt1 release on individual DNAs. We combined data from the first and second Mcm2-7 associations in 57 double-hexamer formation events, and observed 99 Cdt1 association and release events (Figure 4c, Figure 4–figure supplement 3). In the majority of these events (85/99), Cdt1 release is within experimental time resolution of the loss of OM-FRET (Figure 4d). In a smaller number of events (12/99), Cdt1 releases either immediately before or after OM-FRET ends (within ± 7.2s). We confirmed the loss of half the acceptor fluorescence cannot be consistently explained by photobleaching of the acceptor fluorophores on either Mcm2-7^2C-650^ or Cdt1^N-649^ (Figure 4– figure supplement 2). The distribution of intervals separating the end of the high *E*_FRET_ state relative to the corresponding Cdt1 release is centered around 0s (Figure 4d). This is true for both the first and the second Cdt1 release events (Figure 4–figure supplement 4). Additionally, *E*_FRET_ values are high in the 10 s interval before Cdt1 release, and low in the 10 s interval after Cdt1 release (Figure 4–figure supplement 4). Together, these data support a model in which Cdt1 release and the end of OM interaction are simultaneous or near-simultaneous, consistent with a connection between these events.

To test the hypothesis that Cdt1 release is coupled to the end of OM interaction, we monitored OM interaction in a mutant that is defective for Cdt1 release. We purified a version of the Mcm2-7^2C-649^ protein with a mutation in the Mcm5-Mcm3 ATPase active site (Mcm5-R549A or 5RA). The Mcm2-7^5RA^ mutant helicase is defective in Cdt1 release and helicase loading (Coster et al., 2014; Kang et al., 2014; Ticau et al., 2017). In reactions with ORC^5C-549^ we observed associations of single Mcm2-7^2C-649,5RA^ proteins (Ticau et al., 2017). Compared to wild-type Mcm2-7^2C-649^ helicase, OM-FRET that began with Mcm2-7 arrival continued for much longer in the 5RA mutant helicase (average OM-FRET duration was 156 ± 14 s; Figure 4e). In all cases (128/128), we observed OM-FRET for the entire duration that both ORC^5C-549^ (donor) and Mcm2-7^2C-649,5RA^ (acceptor) were present, consistent with its duration being limited by photobleaching or loss of both proteins due to sliding off the DNA end. In contrast, in reactions with wild-type Mcm2-7^2C-649^, the first OM interaction ends soon after the first Mcm2-7 arrival (mean 30 ± 2 s). Thus, the results from the Mcm2-7^5RA^ mutant are consistent with the hypothesis that Cdt1 release is correlated with the end of OM interaction.

### Monitoring Mcm2-7-ORC interactions at a second interface

To investigate the protein movements that allow a single ORC to interact with both helicases, as predicted by the one-ORC (Figure 3a) and ORC-flip (Figure 2a) models, we monitored a second interface between Mcm2-7 and ORC during helicase loading. A recent structural study found evidence for the existence of a helicase-loading intermediate in which ORC interacts with the N-terminal face of Mcm2-7 (Miller et al., 2019). The authors referred to this structure as the “MO complex” because of the opposite orientation of the ORC and Mcm2-7 proteins relative to their orientation in the <u>O</u>CC<u>M</u> structure that we have monitored up to this point (Figure 5a). To ask whether the MO interaction is formed during helicase loading, we modified ORC and Mcm2-7 at sites that are proximal (∼40 Å apart) in the MO structure (Figure 5b) but are much further apart (∼150 Å) in the OCCM (i.e., during the OM interaction studied in Figures 1-4). Specifically, ORC was labeled with a donor fluorophore at the Orc6 C-terminus (ORC^6C-549^) and Mcm2-7 was labeled with an acceptor fluorophore at the Mcm3 N-terminus (Mcm2-7^3N-650^). These proteins function equivalently to their unlabeled counterparts in ensemble helicase-loading experiments (Figure 5–figure supplement 1).

**Figure 5.**
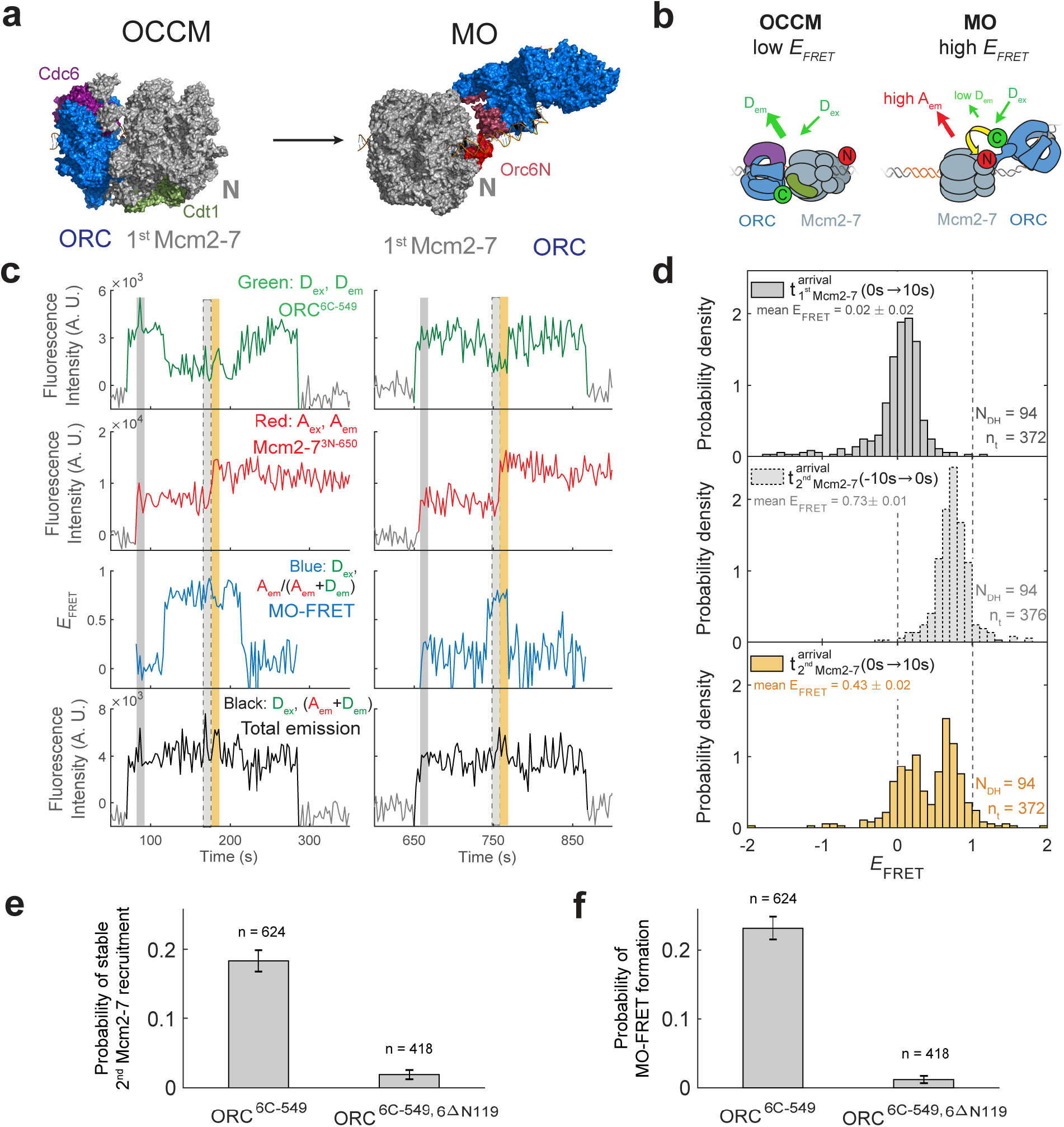
ORC forms a new “MO” interaction interface with the first Mcm2-7 before recruiting the second Mcm2-7 helicase. **(a)** Cryo-EM structures of the <u>O</u>CC<u>M</u> (OM) and MO (PDB: 5V8F and 6RQC) show the inversion of ORC relative to the first Mcm2-7 helicase. **(b)** Schematic of ORC^6C-549^ and Mcm2-7^3N-650^ proteins in the context of the OCCM (left) and MO (right) structures. We predict that we would observe substantial FRET only in the MO but not the OCCM structure. **(c)** Representative traces showing ORC and Mcm2-7 associations with DNA and MO interaction *E*_FRET_ during helicase loading. Donor-excited fluorescence record shows ORC^6C-549^ association (Green: D_ex_, D_em_ panel) and acceptor-excited fluorescence record shows Mcm2-7^3N-650^ association (Red: A_ex_, A_em_ panel). Formation of the MO interaction brings the donor and acceptor fluorophores in proximity resulting in increased FRET efficiency (*E*_FRET_, Blue: D_ex_, A_em_/ (A_em_ + D_em_) panel). The black: D_ex_, (A_em_ + D_em_) panel shows donor-excited total emission. Gray, gray-dash, and yellow highlight the three 10 second (s) time intervals referenced in (d). Concentrations of proteins in the reaction are 0.5 nM ORC^6C-549^, 15 nM Mcm2-7^3N-650^ and 20 nM unlabeled Cdt1. Figure 5–figure supplement 2 shows additional records. **(d)** Histograms of MO interaction *E*_FRET_ values during three 10 s time intervals during helicase loading: 10 s after the first (top) and second (bottom) Mcm2-7 arrived and 10 s before the second Mcm2-7 arrived (middle). Examples of these intervals corresponding to the region highlighted in gray, yellow, and gray-dash in (c). The two dashed lines indicate *E*_FRET_ values of 0 and 1. N_DH,_ number of double-hexamer formation events and n_t_, number of signal points. *E*_FRET_ values below -2 and above +2 were excluded (4/376, 0/376 and 4/376 signal points from the top, middle and bottom histograms resp.). **(e)** The N-terminal domain of ORC6 is required for second Mcm2-7 recruitment. The fraction (± SE) of n first Mcm2-7^3N-650^ binding events that resulted in stable recruitment of a second Mcm2-7^3N-650^ is plotted for WT-ORC^6C-549^ and ORC^6C-549,6ΔN119^ proteins **(f)** The N-terminal domain of ORC6 is requited for MO interaction. The fraction (± SE) of n first Mcm2-7^3N-650^ binding events that resulted in MO-FRET formation is plotted for WT-ORC^6C-549^ and ORC^6C-549,6ΔN119^ proteins.

When ORC^6C-549^ and Mcm2-7^3N-650^ both co-localized with DNA, we observed periods of high *E*_FRET_ (Figure 5c, Figure 5–figure supplement 2; Blue: D_ex_, A_em_/(A_em+_D_em_) panel). Because we labeled the MO interface, we will refer to periods of high *E*_FRET_ arising from this set of probes as “MO interactions”. In almost every observed double-hexamer formation event, we observed a period of MO interaction (94/95 events). These MO interactions occurred only once during helicase loading, and formation of the MO high *E*_FRET_ state (MO-FRET) always anticipated the arrival of the second Mcm2-7 (94/94 events). Unlike the OM interactions we monitored previously, MO interactions do not begin simultaneously with the arrival of the first or second Mcm2-7. Instead, MO *E*_FRET_ in the 10 s period immediately after first Mcm2-7 arrival was low, with an average value of 0.02 ± 0.02 (Figure 5d; top panel). In contrast, *E*_FRET_ values in the 10 s interval just before second Mcm2-7 arrival were high with an average value of 0.73 ± 0.01 (Figure 5d; middle panel). Interestingly, the MO interaction is lost rapidly after the second Mcm2-7 associates with DNA (average MO-FRET duration after second Mcm2-7 arrival was 9.7 ± 1.5 s). This short duration is reflected in the bimodal distribution of the *E*_FRET_ values in the 10 s interval just after second Mcm2-7 arrival (Figure 5d; bottom panel).

To verify that the increases in *E*_FRET_ that we refer to as MO-FRET represent formation of the previously described “MO complex”, we performed experiments with a mutant in Orc6 that disrupts MO formation (Miller et al., 2019). We purified a version of the ORC^6C-549^ protein in which the N-terminal domain of Orc6 was deleted (ORC^6C-549,6ΔN119^). Ensemble helicase loading assays using this protein show a strong reduction in successful helicase loading (Miller et al., 2019). In single-molecule reactions with ORC^6C-549,6ΔN119^ and Mcm2-7^3N-650^, we nearly always observed association of a single Mcm2-7 helicase; only 8/418 first Mcm2-7 association events go on to recruit a stable second Mcm2-7 (Figure 5e). Furthermore, ORC^6C-549,6ΔN119^ showed very few instances with MO-FRET (5/418 of first Mcm2-7 association events; Figure 5f). Both of these frequencies are much lower than what is observed with wild-type ORC^6C-549^ (Figure 5e and 5f). These results are consistent with the MO-FRET signal arising due to formation of the MO complex, and with formation of the MO interaction being critical for second Mcm2-7 recruitment.

Establishment of the MO interaction is likely to be a limiting step in helicase loading. Only a fraction (0.18 ± 0.02) of first Mcm2-7 binding events detected with wild-type ORC convert to stable second Mcm2-7 recruitment (Figure 5e, ORC^6C-549^). Strikingly, this frequency is similar to the fraction of first Mcm2-7 binding events that have MO-FRET (0.23 ± 0.02; Figure 5f, ORC^6C-549^). Consistent with the hypothesis that formation of the MO interaction is the step at which most unsuccessful helicase loading attempts fail, formation of MO-FRET is a strong predictor of stable second Mcm2-7 association; 79 ± 3% of first Mcm2-7 association events with MO-FRET go on to recruit a stable second Mcm2-7.

The likely reason that MO formation is important for successful helicase loading is that the MO complex stabilizes the first Mcm2-7 on the DNA. Consistent with this hypothesis, we find that when MO formation is inhibited in the context of the ORC^6C-549,6ΔN119^ mutant, the first recruited Mcm2-7 dissociates rapidly after ORC is released from the DNA (Figure 5–figure supplement 3a). Under this condition, half the first Mcm2-7 complexes recruited are released in ∼ 11 s of ORC release (Figure 5–figure supplement 3b). This is not simply a consequence of the mutant ORC. Mcm2-7 complexes recruited by wild-type ORC that do not form an MO complex show the same rapid release from DNA after ORC dissociation (Figure 5–figure supplement 3c). In contrast, Mcm2-7s that form the MO interaction have longer dwell-times on DNA (lower limit is ∼ 35 s for loss of half the Mcm2-7 complexes; Figure 5–figure supplement 3d). We note that for the majority of these events the end of the MO interactions occurred as a consequence of second Mcm2-7 recruitment rather than their release from the DNA, indicating that the measured times are underestimated.

### Cdt1 release coordinates the ORC flip on DNA

To examine more directly the order of events surrounding MO formation, we monitored Cdt1 release and MO-FRET in a single experiment. The SM helicase-loading experiment was performed with the MO-FRET pair ORC^6C-549^ and Mcm2-7^3N-650^. Additionally, Cdt1 was labeled at its C-terminus with an acceptor fluorophore (Cdt1^C-650^). Similar to the approach in Figure 4b, association of an Mcm2-7^3N-650^/ Cdt1^C-650^ results in the association of two red fluorophores (Figure 6a, Red: A_ex_, A_em_ panel). The subsequent loss of half the acceptor fluorescence intensity soon after Mcm2-7/Cdt1 arrival is indicative of Cdt1 release. Proximity between donor-labeled ORC^6C-549^ and acceptor-labeled Mcm2-7^3N-650^ results in one period of increased *E*_FRET_ (MO interaction) that starts before the recruitment of the second Mcm2-7. Although Cdt1^C-650^ is also labeled with an acceptor fluorophore, ORC^6C-549^ exhibits very low FRET with Cdt1^C-650^ (average *E*_FRET_ is 0.10 ± 0.01; Figure 6–figure supplement 1). Thus, we can monitor both MO interactions and Cdt1 arrival and departure in these experiments.

**Figure 6.**
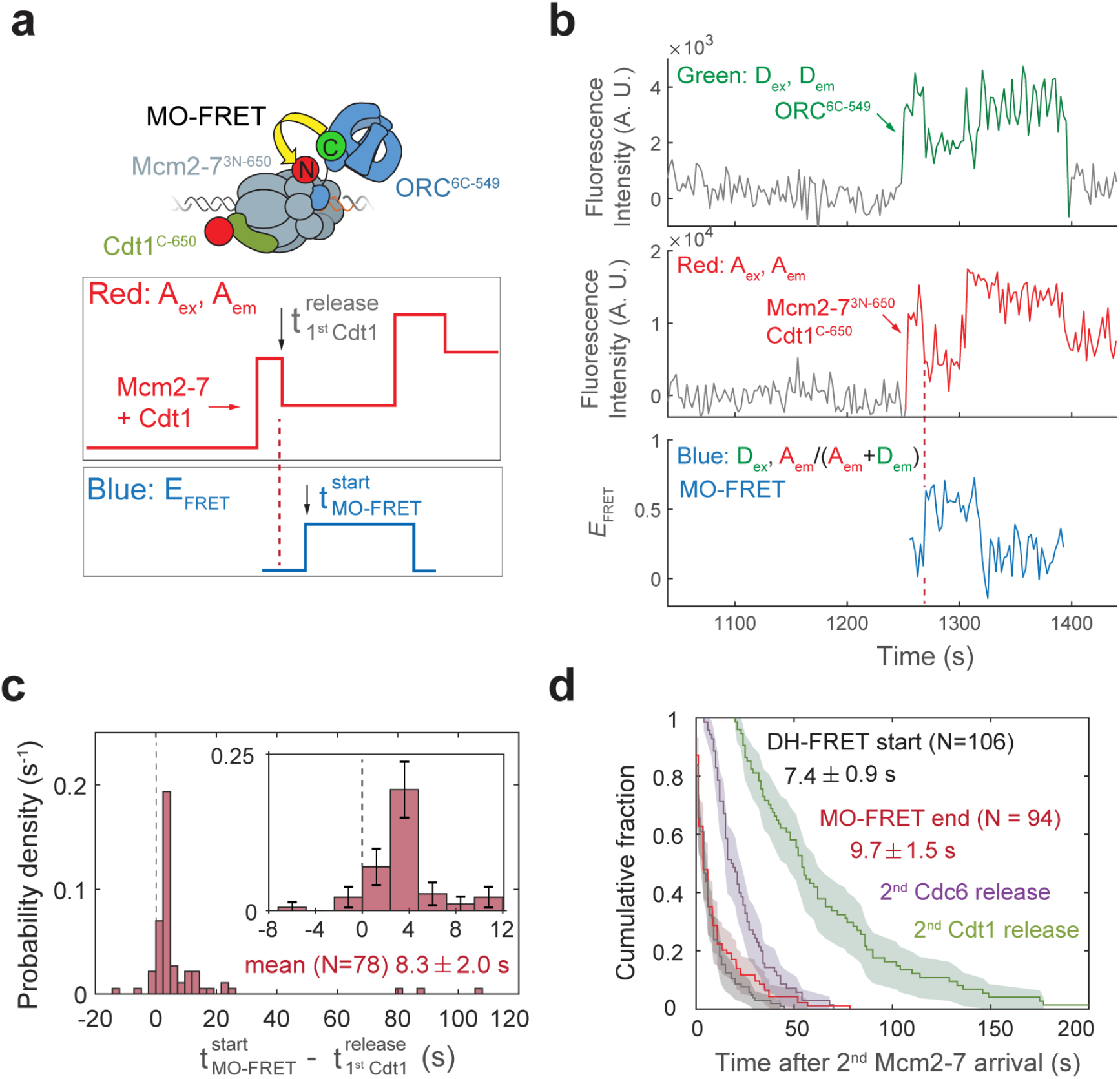
Cdt1 release occurs before MO formation. **(a)** Experimental setup for observation of Cdt1 release and MO interactions on the same DNA molecule. ORC and Mcm2-7 are labeled using the MO-FRET pair ORC^6C-549^ (donor) and Mcm2-7^3N-650^ (acceptor), respectively. Cdt1^C-650^ is also labeled with an acceptor fluorophore, but at a position that exhibits low *E*_FRET_ with ORC^6C-549^ (Figure 6–figure supplement 1). The panels below show the predicted traces if Cdt1 release occurs before MO formation during a helicase-loading event. Each Mcm2-7^3N-650^/Cdt1^C-650^ association results in an increase in fluorescence in the acceptor-excited fluorescence record (Red: A_ex_, A_em_ panel). The black arrow indicates the release of Cdt1^C-650^ that results in a decrease in acceptor-excited fluorescence roughly half the size of the prior increase. **(b)** A representative trace showing the sequential association of two Mcm2-7^3N-650^/Cdt1^C-650^ molecules and MO-FRET. The red dashed line indicates the first Cdt1 release time. Figure 6–figure supplement 3a shows additional records. Concentrations of proteins in the reaction are 0.5 nM ORC^6C-549^, 10 nM Mcm2-7^3N-650^ and 20 nM Cdt1^C-650^. **(c)** Release of the first Cdt1 anticipates the onset of MO interactions. The start-time of MO-FRET after the first Cdt1 release is plotted as a histogram with bin size of 2.4 s. Dashed line indicates a time difference of zero between first Cdt1 release and MO-FRET start time. Inset: magnified view (±SE). **(d)** MO interactions end rapidly after arrival of the second Mcm2-7 with a timing that matches that of double-hexamer formation. Cumulative distributions of four events after second Mcm2-7 arrival are plotted; the time to end of MO interaction (red), start of double-hexamer interactions detected via FRET (Ticau et al., 2015), and release of the second set of Cdc6 and Cdt1 molecules (Ticau et al., 2015). Shaded areas represent the 95% confidence intervals for each curve.

Kinetic analysis of MO-FRET onset indicates that the MO-intermediate is formed shortly after the first Cdt1 is released and the OM-interaction is lost. We analyzed 78 events with stable second Mcm2-7 recruitment and detectable first Cdt1 release (e.g., Figure 6b). We confirmed that the loss of half the acceptor-excited acceptor fluorescence, which we infer as Cdt1 release, is unlikely to be due to photobleaching of the acceptor fluorophore on Cdt1^C-650^ (Figure 6–figure supplement 2). In 76/78 events, MO-FRET is either established simultaneously (within ± 2.4 s) with release of the first Cdt1 or occurs after the first Cdt1 is released. Strikingly, MO interactions began after Cdt1 release in a majority of these events (59/78 events; Figure 6c). The average time to MO-FRET after first Cdt1 release is 8.3 ± 2.0 s indicating that establishment of MO-FRET is a separate, subsequent event from the end of the first OM-FRET, which is simultaneous with the first Cdt1 release (Figure 4d). Consistent with the first Cdt1 release occurring prior to formation of the MO interaction, MO-*E*_FRET_ values are mostly low in periods when the first Cdt1 is associated and high after the first Cdt1 is released (Figure 6–figure supplement 3).

To further establish the order of first Cdt1 release and formation of the MO interaction, we monitored the formation of MO interactions in a mutant that is defective for Cdt1 release. To this end we used the same Mcm2-7^5RA^ mutant that is defective for Cdt1 release and second Mcm2-7 recruitment but now labeled to detect MO interactions. In reactions with ORC^6C-549^ and Mcm2-7^3N-650,5RA^ molecules, we observed only single Mcm2-7 association in 162/163 Mcm2-7 binding events (Figure 6–figure supplement 4a). Consistent with Cdt1 release being required for MO formation, only 3/163 events showed MO interactions (fraction of events is 0.02 ± 0.01). This is a much lower fraction than is observed for the wild-type protein (0.23 ± 0.02, Figure 6–figure supplement 4b). Together, these data support a model in which Cdt1 release anticipates formation of the MO interaction.

What drives the dissolution of the MO complex? There are three temporally separable events known to occur after arrival of the second Mcm2-7: 1) initiation of double-hexamer interactions between the two helicases (Ticau et al., 2015); 2) release of the second Cdc6 (Ticau et al., 2015); and 3) three typically simultaneous events of release of the second Cdt1, closing of the second Mcm2-7 ring, and release of ORC (Ticau et al., 2017). We compared the distribution of times of loss of MO interaction relative to the arrival of the second Mcm2-7 to the previously determined times of each of these events (Figure 6d). The distribution of second Cdc6 and second Cdt1 release times are clearly distinct from the times of MO dissolution. In contrast, we observed a similar distribution for the end of the MO interaction after arrival of the second Mcm2-7 (Fig. 6e, 9.7 ± 1.5 s) relative to the time required for initial double-hexamer interactions after second Mcm2-7 arrival (Ticau et al., 2015). These data strongly suggest that formation of double-hexamer interactions between the first and second helicase accompany a structural reconfiguration that ends the MO interaction.

## Discussion

By monitoring interactions between ORC and Mcm2-7 through two separate interfaces, the single-molecule experiments described here provide unique insights into helicase loading. We demonstrate that similar, if not identical, interactions between DNA-bound ORC and the C-terminal regions of Mcm2-7 occur during recruitment of the two helicases during double-hexamer formation (OM interactions, Figures 1 and 2). Importantly, we show that one ORC is sufficient to efficiently recruit both helicases (Figure 3) but that in rare cases separate ORC molecules may recruit the two helicases. Between the two helicase recruitment events, ORC forms a distinct interface with the N-terminal region of the first recruited Mcm2-7 (MO interaction, Figure 5) that mediates retention of ORC on the DNA and is required for recruitment of the second Mcm2-7. Finally, our experiments define the order for Cdt1 release and establishment of double-hexamer interactions relative to OM and MO interactions (Figures 4 and 6). Overall our findings reveal a highly coordinated series of events ensuring the formation of the correct head-to-head conformation of the loaded helicases.

### A single-ORC model for helicase loading

Based on our findings, we propose a detailed model for helicase loading by one ORC (Figure 7). ORC/Cdc6 bound to origin DNA recruits the first Cdt1-bound-Mcm2-7 helicase via the first OM interaction (Figure 7; OC^6^C^1^M_1_ state). This complex is maintained until the ATP-hydrolysis dependent releases of Cdc6 and then Cdt1, forming the OM_1_ state. Following release of the first Cdt1, ORC releases from its initial DNA and Mcm2-7 binding sites to form a distinct interaction with the N-terminal region of the first Mcm2-7 (pre-M_1_O state). This tethered conformation prevents ORC from releasing into solution as it exchanges DNA binding sites. As a consequence of the ORC inversion, the two helicases are recruited in opposite orientations (M_1_OC^6^C^1^M_2_ state) and rapidly interact to form a head-to-head Mcm2-7 double hexamer (M_1_M_2_).

**Figure 7.**
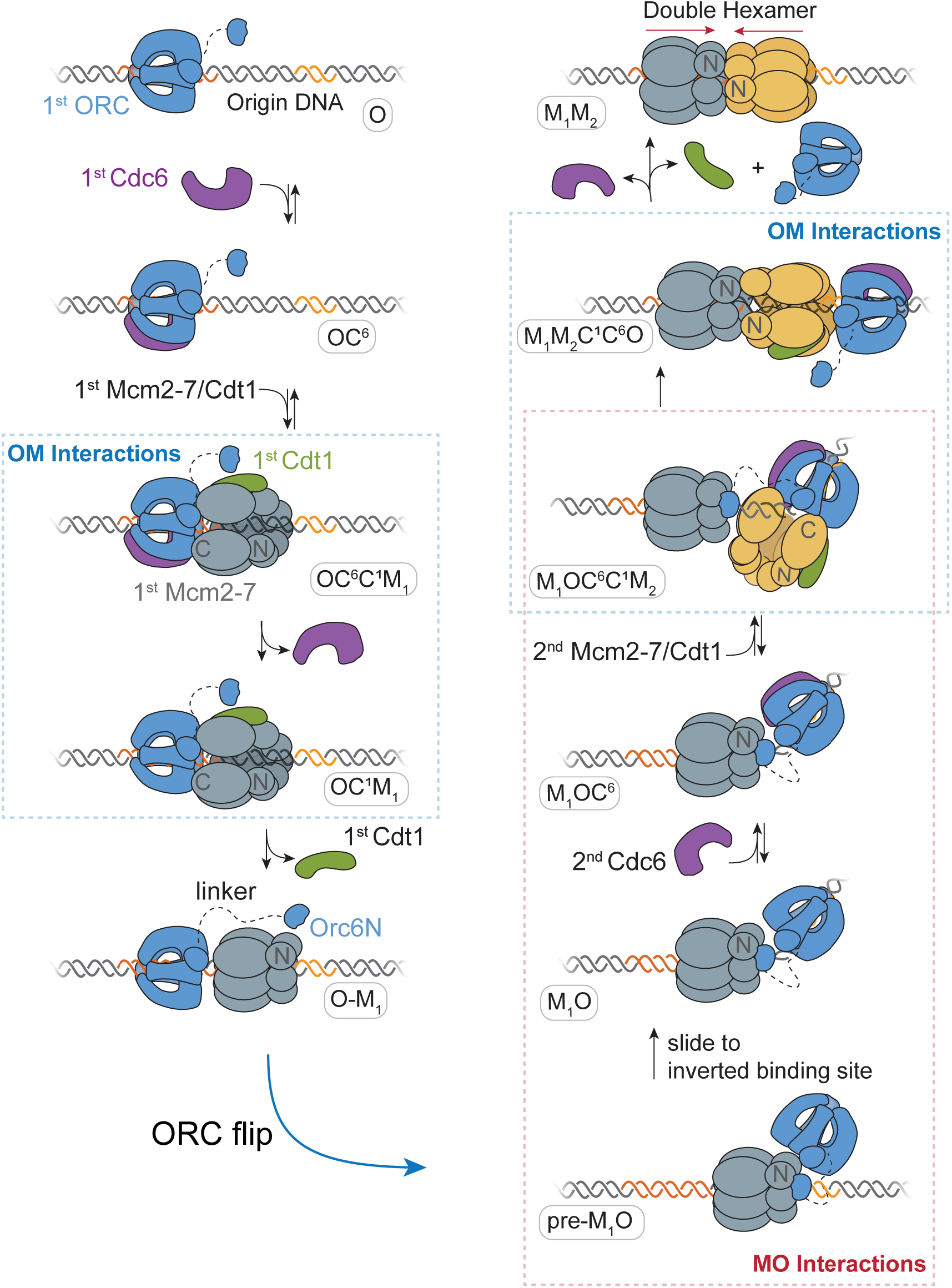
A model for helicase loading mediated by one ORC molecule. Designations in ovals are unique names for each intermediate using the abbreviations O, ORC; C^6^, Cdc6; C^1^, Cdt1; M_1_, 1^st^ Mcm2-7; and M_2_, 2^nd^ Mcm2-7. Dashed curve represents the linker that connects the N- and C-terminal domains of Orc6. Orange and yellow DNA segments are sequence-specific ORC binding sites, see text for details. Note that the ORC and Mcm2-7 proteins are not interacting in the O-M_1_ complex.

The model we present reconciles previous observations that seemed contradictory. A single ORC making similar OM interactions during recruitment of both Mcm2-7 complexes is consistent with previous observations that Mcm2-7 mutations in this interface interfere with recruitment of both the first and second helicase (Coster and Diffley, 2017; Frigola et al., 2013). Further, the proposal that ORC recruits both helicases by re-binding DNA in an inverted orientation corresponds with a requirement for two ORC-binding sites at origins (Coster and Diffley, 2017). We show that the MO interaction is consistently present during recruitment of the second Mcm2-7. This finding is consistent with structural studies suggesting that an ORC molecule engaged with the N-terminal region of the first Mcm2-7 is responsible for recruiting the second Mcm2-7 (Miller et al., 2019). Because the MO complex holds ORC in close proximity to the first Mcm2-7 as the second Mcm2-7 is recruited, this finding explains the rapid formation of double-hexamer interactions after second Mcm2-7 arrival (Ticau et al., 2015). Finally, our observation that a single ORC molecule can guide helicase loading is consistent with previous single-molecule studies and time-resolved cryo-EM studies, both of which observed predominantly one ORC engaged with DNA during helicase loading (Miller et al., 2019; Ticau et al., 2015).

Our experiments reveal that the events in helicase loading guided by a single ORC are highly coordinated. The first OM interaction begins simultaneously with arrival of the first Mcm2-7/Cdt1 and ends when the first Cdt1 is released. The MO interaction is formed shortly after the first Cdt1 is released and the OM interaction ends (mean 8.7 ± 2.0 s after Cdt1 release), and anticipates recruitment of the second Mcm2-7. Consistent with this ordering, preventing Cdt1 release with a Mcm2-7 ATPase mutant extends the duration of OM interactions (average OM duration after first Mcm2-7 arrival is 30 ± 2 s for wild-type and 156 ± 14 s for the Mcm2-7^5RA^ mutant) and prevents formation of MO interactions. The second OM interaction begins simultaneously with the second Mcm2-7 arrival (M_1_OC^6^C^1^M_2_ state). We infer that establishment of the MO interaction is required to recruit the second Mcm2-7 helicase based on two observations. First, we consistently observe that the onset of the MO interaction anticipates recruitment of the second Mcm2-7. Second, disrupting the MO interaction using a mutant version of ORC prevents second Mcm2-7 recruitment (Figure 5). Finally, the temporal connection between the formation of double-hexamer interactions and the end of the MO interactions (Figure 6d), strongly suggests that formation of double-hexamer interactions breaks the MO interaction. This supports the existence of two complexes with the same set of proteins but distinct ORC-Mcm2-7 and Mcm2-7-Mcm2-7 interactions (Figure 7, M_1_OC^6^C^1^M_2_ and M_1_M_2_C^1^C^6^O states). The requirement to form the MO complex to recruit the second Mcm2-7 combined with the requirement to disrupt the MO complex to establish double-hexamer interactions together provide an ordered selection to ensure that successfully recruited second Mcm2-7 helicases go on to form double-hexamers.

We propose that a single ORC molecule uses a tether to the first helicase to remain DNA associated while ORC locates and binds a second inverted DNA site. This mechanism is supported by the observation that ORC is not bound to the primary ORC-binding site on DNA in a structure of the MO complex (Miller et al., 2019). Although protein flipping has not been reported in other steps of DNA replication, multiple DNA-binding proteins have been shown to switch orientations on DNA (Ganji et al., 2016). In these previously reported systems, protein inversion is proposed to occur via dissociation into solution, followed by rapid reorientation and rebinding resulting in the flipped protein binding the same (quasi-)palindromic DNA site (Marklund et al., 2020; Sasnauskas et al., 2011), or a closely-spaced inverted binding site (Ryu et al., 2014). The ORC gymnastics observed here however are distinguished from other protein flipping events by the use of a protein tether between ORC and the first Mcm2-7 helicase.

### Origin structure and helicase loading

A model in which the MO complex is required to recruit the second Mcm2-7 is consistent with origin architecture in budding yeast, wherein two ORC-binding sites with different affinities are required for helicase loading *in vivo*. Previous studies have shown that in addition to the primary ORC-binding site (<u>A</u>RS <u>c</u>onsensus <u>s</u>equence or ACS), natural origins consistently have at least one (and sometimes multiple) additional weak, inverted ORC-binding sites referred to as B2 elements (Bell and Stillman, 1992; Marahrens and Stillman, 1992; Palzkill and Newlon, 1988; Rao et al., 1994; Wilmes and Bell, 2002). Interestingly, even though two oppositely-oriented strong ORC-binding sites support origin function both *in vitro* and *in vivo* (Coster and Diffley, 2017), natural origins have selected against this architecture (Chang et al., 2011; Marahrens and Stillman, 1992; Palzkill and Newlon, 1988; Rao et al., 1994). This selection argues against a model in which ORC is pre-bound at both sites. Indeed, when B2 elements have been mapped they are frequently located at positions in which the presence of the OCCM (OC^6^C^1^M_1_ in Figure 7) would clash with ORC binding at B2 (Chang et al., 2011; Marahrens and Stillman, 1992; Rao et al., 1994). Along these lines, in the small number of two-ORC events in our studies, we observe second ORC associations occur after the first, but before the second, Mcm2-7 arrives. Thus, we propose that, whether one or two ORCs mediate loading, the ORC mediating second Mcm2-7 recruitment would only stably associate with B2 after the first Mcm2-7 is recruited. Such a mechanism would ensure that these sites are only bound once the first Mcm2-7 is recruited, thus driving the reaction towards double-hexamer formation and preventing two ORC proteins independently recruiting “first” Mcm2-7 complexes.

The relative placement of the two ORC-binding sites might influence the coordinated series of events we observe in helicase loading. Although the two oppositely-oriented ORC-binding sites are very close in the *ARS1* origin used in our experiments, the distance between these sites is variable across natural origins (Chang et al., 2011; Palzkill and Newlon, 1988). We speculate that the pre-M_1_O slides in search of a second, inverted ORC-binding site (Figure 7) and recruits the second Mcm2-7 only once an inverted ORC-binding site is found. As described above, for a number of origins for which B2 elements have been identified (including *ARS1*) this sliding would be required to expose the B2 element (in Figure 7, sliding to the left). A number of helicase-loading intermediates slide *in vitro* (Sánchez et al., 2021; Scherr et al., 2021), and experiments placing barriers to sliding at *ARS1* support a role for sliding to identify second inverted ORC-binding sites (Coster and Diffley, 2017; Warner et al., 2017). Consistent with the argument that second Mcm2-7 recruitment only occurs when the sliding pre-M_1_O finds a second inverted ORC-binding site, helicase loading does not occur at artificial origins with a single strong ORC-binding site (Coster and Diffley, 2017).

### Comparison of one-ORC and two-ORC mechanisms for helicase loading

Our data also demonstrate that two distinct ORC molecules are present in a small subset of helicase loading events and provide information about the mechanism of these events. One insight comes from the analysis of first Mcm2-7 DNA retention in the absence of MO formation. Analyses of helicase loading with the ORC^6ΔN119^ mutant and first Mcm2-7 helicases that fail to form the MO interaction show that in both cases the Mcm2-7 molecules are rapidly released from the DNA after ORC release (Figure 5–figure supplement 3). This finding indicates that the first Mcm2-7 is unstable if the MO complex is not formed and that there would be only a short time window for a second ORC molecule to form an MO complex before the first Mcm2-7 is released. It is also interesting to consider how the MO complex is formed by a second ORC. There are two possible ways for a second ORC to interact with the first Mcm2-7 (Figure 7–figure supplement 1). It is possible that the second ORC is recruited out of solution by MO-like interactions with the first Mcm2-7 and then identifies an appropriate B2 DNA binding site. Alternatively, ORC could bind at the low-affinity B2 site first and then form the MO-interaction with the first Mcm2-7. In both cases, there would be a short time window for the second ORC to form the MO before the first Mcm2-7 is released. Analysis of the two-ORC events in the assay that allows detection of the MO complex suggests both mechanisms are used.

It is likely that both one-ORC or two-ORC mechanisms are used *in vivo* and it will be difficult to determine which mechanism is used more frequently. The low frequency of two-ORC events we observe (see Figure 3) suggests that the first ORC has a higher probability of forming the MO complex through a helicase-tethered flip than a second ORC, consistent with the first ORC already being associated with the first Mcm2-7. On the other hand, our single-molecule experiments are necessarily performed at low ORC concentrations. Given the short time window for a second ORC to act, higher ORC concentrations would be predicted to increase the percentage of two-ORC events. Finally, we note that it is difficult to determine the free ORC concentration *in vivo,* particularly given the ability of ORC to interact non-specifically with AT-rich DNA. Nevertheless, consistent with ORC being limiting in cells, *in vivo* foot-printing studies show that only a subset of the origin-associated ACSs are strongly occupied in cells (Belsky et al., 2015). This finding suggests it is unlikely that ORC spends significant time bound to the much weaker B2 ORC binding sites at origins.

### Cdt1 release coordinates the transition between OM and MO complexes

Our data shows that release of the first Cdt1 is a critical regulatory step in helicase loading. There are two potential explanations for the simultaneous end of OM interactions and Cdt1 release observed here. Cdt1 binding could be required for OM interactions such that Cdt1 release reduces or eliminates these interactions. Consistent with this hypothesis, Cdt1 binding has been proposed to alleviate an autoinhibitory function of Mcm2-7, thereby allowing interaction with ORC/Cdc6/DNA (Fernández-Cid et al., 2013). A second possibility is that closure of the Mcm2-5 gate, which is concomitant with Cdt1 release (Ticau et al., 2017), induces a conformational change in the Mcm2-7 complex that disrupts the OM interaction. It is intriguing to note that ORC is frequently released from the DNA simultaneously with the second Cdt1 release (Ticau et al., 2015), suggesting that in addition to breaking the OM interaction, release of the first Cdt1 may stimulate ORC release from DNA.

Release of the first Cdt1 and the concomitant closure of the Mcm2-5 gate around DNA likely create a new interaction site on Mcm2-7 for Orc6 (Miller et al., 2019; Ticau et al., 2017). The N- and C-terminal domains of Orc6 (Orc6N and Orc6C) interact with Cdt1 *in vitro* (Chen and Bell, 2011). We propose that once the first Cdt1 is released, Orc6N is able to reach over the first Mcm2-7 and establish a tether with the Mcm2 N-terminal region (Miller et al., 2019). In this model, the extensive linker (∼160 amino acids) between the Orc6N and Orc6C serves as flexible leash that facilitates a search for this new Orc6N binding site (Chen and Bell, 2011). Since ORC is not Cdc6-bound at this stage (OM_1_ state), ORC forms an open ring around DNA and can release from its primary DNA-binding site. Additional events that could contribute to ORC release from its initial high-affinity binding site, include the conformational changes driving dissolution of the OM complex and changes in ORC conformation upon binding to the N-terminal region of Mcm2-7. The closed Mcm2-5 gate in the M_1_O facilitates contacts between Orc6C and the Mcm5 N-terminal region such that the two domains of Orc6 form a latch across the Mcm2-5 gate (Miller et al., 2019), a likely mechanism for the MO complex stabilizing DNA association of the first Mcm2-7 after loss of the OM interaction (Figure 5–figure supplement 3). We show that Orc6N is required to form the MO interaction, and that release of the first Cdt1 and formation of the MO interaction occur in close succession. However, we could not determine the timing of establishment of the Orc6N tether (as in the pre-M_1_O intermediate) relative to the end of the first OM interaction (or the first Cdt1 release), as fluorophore modification of Orc6 N-terminal domain interfered with helicase loading.

### Implications for origin licensing at non-sequence defined origins

A single-ORC mechanism for helicase loading has potential implications for DNA replication initiation in higher eukaryotes. Although the helicase-loading proteins from budding yeast are highly conserved, ORC does not exhibit sequence-specific binding in higher eukaryotes where origins are not defined by specific DNA sequences (Bleichert et al., 2017; Prioleau and MacAlpine, 2016; Vashee et al., 2003). A mechanism in which two separate ORC molecules guide loading of the two helicases would require two independent non-specific ORC-binding events. Each single Mcm2-7 helicase would then have to find an oppositely-oriented helicase to complete helicase loading. In contrast, helicase loading guided by a single ORC that performs an inversion places fewer constraints on where helicases can be loaded and could still be supported with low ORC concentrations. Additionally, a full round of helicase loading completed by a single ORC could prevent non-productive loading of single helicases that cannot support bidirectional replication (Champasa et al., 2019; Miller et al., 2019).

## Methods

### Protein purification and labeling

Wild-type ORC and Cdc6 were purified as described previously (Frigola et al., 2013). We used a N- and C-terminal protein modifications to fluorescently label ORC (-LPETGG at the C-terminus of Orc5 and Orc6, Ubiquitin-GGG-Flag at the N-terminus of Orc1), Mcm2-7 (LPETGG at the C-terminus of Mcm2, Ubiquitin-GGG-Flag at the N-terminus of Mcm4, 3xFlag-TEV[ENLYFQ/G]-GG at the N-terminus of Mcm3) and Cdt1 (Ubiquitin-GGG tag at the N-terminus, LPETGG tag at the C-terminus). The ubiquitin (*in vivo*) and 3xFlag-TEV (using TEV protease, NEB) fusions were removed to reveal three N-terminal glycines required for N-terminal sortase-mediated labeling. The peptides NH_2_-CHHHHHHHHHLPETGG-COOH and NH_2_GGGHHHHHHHHHHC-COOH were used for N- and C-terminal labeling respectively and will be referred to as the N-peptide and C-peptide. The N- and C-peptides were labeled with maleimide-derivatized DY549P1 (Dyomics), DY649P1 (Dyomics), Dylight550 (Thermo-scientific), Dylight650 (Thermo-scientific), or Cy3B (Cytiva) via cystine-maleimide conjugation as described previously (Ticau et al., 2015). Sortase was used to couple the fluorescently-labeled peptide to the N- or C-terminus of the helicase-loading proteins as described below. The peptide-coupled proteins were separated from uncoupled proteins using Ni-NTA Agarose (Qiagen).

### Note on fluorophore-modified protein nomenclature

We use a shorthand for the site of fluorescent modification and the fluorophore to indicate various labeled proteins (for e.g. ORC^5C-549^). The numerical superscripts 549 and 649 indicate proteins coupled to the fluorophores DY549P1 and DY649P1 respectively. Similarly, the superscripts 550 and 650 indicate the fluorophores Dylight550 and Dylight650. DY549P1 and Dylight550 have almost identical spectral properties, and the same is true for DY649P1 and Dylight650. The numerical subscripts are preceded by the site of fluorescent labeling on the protein. Thus, ORC^5C-549^ indicates ORC labeled at the C-terminus of Orc5 with the DY549P1 dye.

### Preparation of labeled ORC^5C-549^, ORC^5C-Cy3B^ and ORC^6C-549^

*S. cerevisiae* (W303 background) strains ySG10 (CBP-Orc1, Orc2-4, Orc^5C-LPETGG^, Orc6) and ySG40 (CBP-Orc1, Orc2-5, Orc^6C-LPETGG^) were grown to OD_600_ = 1.2 in 8 L of YEP supplemented with 2% glycerol (w/v) at 30°C. Cells were arrested in G1 using α-factor (100 ng/mL) and ORC expression was induced using 2% galactose (w/v). After 6 hr, cells were harvested and sequentially washed with 150 mL ORC wash buffer (25 mM Hepes-KOH pH 7.6, 1 M Sorbitol) and ORC lysis buffer (25 mM Hepes-KOH pH 7.6, 0.05% NP-40, 10% Glycerol + 0.5 M KCl). The washed pellet was resuspended in approximately 1/3 of packed cell volume of ORC lysis buffer containing cOmplete Protease Inhibitor Cocktail Tablet (1 tablet per 25 mL total volume; Roche) and frozen dropwise in liquid nitrogen.

Frozen cells were lysed in a SamplePrep freezermill (SPEX) and the lysate was clarified by ultracentrifugation in a Type 70 Ti rotor at 45 krpm for 1 hr at 4°C. The supernatant was supplemented with 2 mM CaCl_2_ and applied to 2.4 mL bed volume (BV) Calmodulin-affinity resin (Agilent) pre-equilibrated in ORC lysis buffer containing 2 mM CaCl_2_. The supernatant and resin were incubated with rotation for 1.5 hr at 4°C. The resin was collected on a column and the flow-through was discarded. The resin was then washed with 10 BV of Buffer A (50 mM Hepes-KOH pH 7.6, 5 mM Mg(OAc)_2_, 10% glycerol) supplemented with 0.2 M KCl, 0.02% NP-40 and 2mM CaCl_2_. ORC was eluted with 5 BV buffer A containing 0.2 M KCl, 0.02% NP-40, 1 mM EDTA and 2 mM EGTA, with 15 min incubations before each elution. Elutions containing ORC (typically fractions 2-4) were pooled and applied to 1 mL BV SP Sepharose Fast Flow (Cytiva) pre-equilibrated with Buffer A containing 0.2 M KCl and 0.02% NP-40. The resin was washed with 10 BV of Buffer A containing 0.2 M KCl and 0.02% NP-40. ORC was eluted with 5 BV of Buffer A containing 500 mM KCl, 0.02% NP-40. Note that for the ORC protein with the Ubiquitin-GGG tag (the N-terminal methionine is replaced with the ubiquitin followed by three glycines), the N-terminal ubiquitin is cleaved off *in vivo* resulting in three glycines at the N terminus of Orc1. Starting with 8 L of cells, the yield was typically 3 mg of ORC.

About 3 nmol of purified ORC (1.5 mg, <0.9 mL) was incubated with equimolar amount of Srt5° evolved sortase (Chen et al., 2011; Ticau et al., 2015) and CaCl_2_ was added to a final concentration of 5 mM. This was mixed with 100 nmol of either C-peptide labeled with DY549-P1 (for ORC^5C-549^ and ORC^6C-549^) or C-peptide labeled with Cy3B (for ORC^5C-Cy3B^). The reaction was incubated at room temperature for 15 min, and then quenched with 20 mM EDTA. After dye-coupling, the reaction was applied to a Superdex 200 10/300 gel filtration column equilibrated in buffer B (50 mM Hepes-KOH pH 7.6, 5 mM Mg(OAc)_2_, 10% glycerol, 0.3 M potassium glutamate (KGlu), 0.02% NP-40) containing 10 mM imidazole. Peak fractions containing peptide-coupled ORC were pooled and incubated with 0.3 mL of Ni-NTA Agarose Resin (Qiagen) pre-equilibrated in buffer B with 10 mM imidazole, for 1.5 hr with rotation at 4°C. The flow-through was discarded and the resin was washed sequentially with 3 mL buffer B with 15 mM imidazole and buffer B with 25 mM imidazole. Peptide-coupled ORC was eluted using buffer B with 0.3 M imidazole. Peak fractions were pooled, aliquoted, and stored at −80°C.

### Preparation of labeled ORC^6C-549,6ΔN119^

The *S. cerevisiae* strain ySG41 (CBP-Orc1, Orc2-5, 3x-Flag-ΔN119-Orc6^C-LPETGG^) was grown to OD_600_ = 1.2 in 12 L of YEP supplemented with 2% glycerol, then induced, harvested, lysed and eluted off a calmodulin affinity column as described above. Fractions containing ORC were pooled and applied to 1 mL BV Anti-M2 Flag resin (Sigma-Aldrich) which was pre-equilibrated with buffer A containing 0.2 M KCl and 0.02% NP-40. The Flag resin was incubated with rotation for 2 hr at 4°C after which it was collected on a column and the flow-through was discarded. The resin was washed with 10 BV of Buffer A containing 0.2 M KCl and 0.02% NP-40. ORC was eluted with 5 BV buffer A containing 0.2 M KCl, 0.02% NP-40 and 0.3 mg/mL 3xFlag peptide with 30 min incubations before each elution. Fractions containing ORC were purified on SP Sepharose Fast Flow, sortase coupled to C-peptide labeled with DY-549P1, and further purified by gel filtration and Ni-NTA columns as described above. Peak fractions were aliquoted and stored at −80°C.

### Preparation of labeled ORC^1N-550^

The *S. cerevisiae* strain yAZ55 (^Ubiquitin-GGG-3xFlag^Orc1, Orc2-6) was grown to OD_600_ = 1.2 in 8 L of YEP supplemented with 2% glycerol, then cell-cycle arrested, induced, harvested, lysate prepared as described above. The lysate was applied to 1 mL BV Anti-M2 Flag resin (Sigma-Aldrich) which was pre-equilibrated with ORC lysis buffer. Flag resin was incubated with rotation for 2 hr at 4°C after which it was collected on a column and the flow-through was discarded. The resin was washed with 10 BV of Buffer A containing 0.2 M KCl and 0.02% NP-40. ORC was eluted with 5 BV buffer A containing 0.2 M KCl, 0.02% NP-40, and 0.3 mg/mL 3xFlag peptide with 30 min incubations before each elution. Fractions containing ORC were pooled and purified on SP Sepharose Fast Flow (Cytiva) as described above. Fractions containing ORC were sortase coupled to N-peptide labeled with Dylight-550, and further purified by gel filtration and Ni-NTA columns as described above. Peak fractions were aliquoted and stored at −80°C.

### Preparation of labeled Mcm2-7^2C-649^ and Mcm2-7^4N-650^

The *S. cerevisiae* strains yST210 (Mcm2^C-LPETGG^, 3xFlag-Mcm3, Mcm4-7, Cdt1) and yST180 (Mcm2, 3xFlag-Mcm3, ^Ubiquitin-GGG^Mcm4, Mcm5-7, Cdt1) were purified as described previously (Ticau et al., 2015, 2017) with the following modifications. We used sortase to couple Mcm2^C-LPETGG^ (yST210) to C-peptide conjugated to DY-649P1 and ^GGG^Mcm4 (yST180) to N-peptide conjugated to Dylight550. Following elution off the Anti-M2 FLAG resin, peptide coupling and gel filtration (Superdex 200 10/300), peak fractions containing peptide-coupled Mcm2-7 were pooled. These pooled fractions were applied to 0.3 mL of Ni-NTA Agarose Resin (Qiagen) pre-equilibrated in buffer B with 10 mM imidazole. The resin was incubated for 1.5 hr with rotation at 4°C. The flow-through was discarded and the resin was washed sequentially with 3 mL buffer B with 25 mM imidazole, 50 mM imidazole and 75 mM imidazole. Peptide-coupled Mcm2-7 was eluted using buffer B with 0.3 M imidazole. Peak fractions were aliquoted, and stored at −80°C.

### Preparation of labeled Mcm2-7^2C-650^, Mcm2-7^2C-650,5RA^

The *S. cerevisiae* strains for Mcm2-7 expression in the absence of Cdt1, ySG12 (Mcm2^C-LPETGG^) and ySG32 (Mcm2^C-LPETGG^, Mcm5RA) were purified in the same manner as the Cdt1/Mcm2-7 complex as described previously (Ticau et al., 2015, 2017). We used sortase to couple Mcm2^C-LPETGG^ (ySG12 and ySG32) to C-peptide conjugated to Dylight650. The Ni-NTA selection for labeled protein was modified as described above for Mcm2-7^2C-649^.

### Preparation of labeled Mcm2-7^3N-650^ and Mcm2-7^3N-650,5RA^

The *S. cerevisiae* strains for Mcm2-7 expression in the absence of Cdt1, ySG24 (^3xFlag-TEV[ENLYFQ/G]-GG^Mcm3), and ySG33 (^3xFlag-TEV[ENLYFQ/G]-GG^Mcm3, Mcm5RA) were grown, harvested and lysate prepared in the same manner as the Cdt1-Mcm2-7 complex as described previously (Ticau et al., 2015). The lysate was applied to 1 mL BV Anti-M2 Flag resin (Sigma-Aldrich) in buffer C (50 mM Hepes-KOH pH 7.6, 5 mM Mg(OAc)_2_, 10% glycerol, 0.3 M KGlu, 0.02% NP-40, 1 mM ATP). The Flag resin was incubated with rotation for 2 hr at 4°C after which it was collected on a column and the flow-through was discarded. The resin was washed with Buffer C and resuspended to create a 50% slurry. ^GGG^Mcm3 (in association with the other Mcm2-7 subunits) was cleaved off the resin with 500 units of 7×His-TEV protease (NEB; rotation overnight at 4°C). The flow-through was collected and applied to 0.4 mL volume Ni-NTA Agarose resin (Qiagen) to remove the 7×His-TEV protease. The flow-through containing cleaved Mcm2-7 (with the three N-terminal glycines revealed) was coupled to N-peptide labeled with Dylight-650 using sortase, and further purified by gel filtration and Ni-NTA columns as described above. Peak fractions were aliquoted and stored at −80°C.

### Preparation of labeled Cdt1^N-649^, Cdt1^C-650^

The *S. cerevisiae* strains for Cdt1 purification yST103 (^Ubiquitin-GGG-^Cdt1-Flag) and ySG046 (3xFlag-Cdt1^C-LPETGG^) were grown to OD_600_ = 1.2 in 8 L of YEP supplemented with 2% glycerol, then cell-cycle arrested, induced, harvested, lysate prepared as described for the Cdt1/Mcm2-7 complex (Ticau et al., 2015). The lysate was applied to 1 mL bed volume (BV) Anti-M2 Flag resin (Sigma-Aldrich) in buffer D (50 mM Hepes-KOH pH 7.6, 5 mM Mg(OAc)_2_, 10% glycerol, 150 mM KGlu, 1mM ATP, 0.02% NP-40) and eluted with buffer D containing 0.3 mg/mL 3xFlag peptide. Fractions containing Cdt1 were sortase coupled to N-peptide labeled with DY649-P1 (for Cdt1^N-649^) or C-peptide labeled with Dylight650 (for Cdt1^C-650^) as described for ORC above. The reaction was applied to a Superdex 75 10/300 gel filtration column equilibrated in buffer D containing 10 mM imidazole. Peak fractions containing peptide-coupled Cdt1 were pooled and incubated with 0.3 mL of Ni-NTA Agarose Resin (Qiagen) pre-equilibrated in buffer D with 10 mM imidazole for 1.5 hr with rotation at 4°C. The flow-through was discarded and the resin was washed sequentially with 3 mL buffer D with 25 mM, 50 mM and 75 mM imidazole. Peptide-coupled Cdt1 was eluted using buffer D with 0.3 M imidazole. Peak fractions were pooled, aliquoted, and stored at −80°C.

### Single-molecule assay for helicase loading

The micro-mirror total internal reflection (TIR) microscope used for single-molecule helicase loading experiments (excitation wavelengths 488, 532, and 633 nm) is described in (Friedman and Gelles, 2012; Friedman et al., 2006). Single-molecule reactions were performed as described in (Ticau et al., 2015) with the following modifications. The surface of the reaction chamber was cleaned by sonication with 2% Micro-90 (Cole Parmer), 0.1M KOH and 100% ethanol sequentially for 1-hr each. The cleaned chamber was then derivatized with a 500:1 ratio of mPEG-silane-2000 (creative PEGWorks) and biotin-mPEG-silane-3400 (Laysan Bio).

Streptavidin-labeled fiducial markers (0.04 micron, Invitrogen TransFluoSpheres) were flowed onto the reaction chamber at a 1:400,000 dilution. A biotinylated 1.3-kb-long origin DNA template (*ARS1*) labeled with Alexa Fluor 488 was coupled to the surface of a reaction chamber using streptavidin. DNA molecules were identified by acquiring two to three images with 488 nm excitation at the start of the experiment. All helicase loading reactions were performed in the reaction buffer as described previously (Kang et al., 2014; Ticau et al., 2015) containing 25 mM Hepes KOH pH 7.6, 7.5 mM Mg(OAc)_2_, 50 μM Zn(OAc)_2_, 100 μM EDTA, 3 mM ATP, 0.3 M KGlu, 12.5 mM DTT, 2.5 mg/mL BSA, an oxygen scavenging system (glucose oxidase/catalase) and triplet state quenchers to minimize photobleaching (Friedman et al., 2006; Hoskins et al., 2011). All reactions additionally contained 1 μM of a 60-bp double-stranded DNA that cannot compete for ORC binding in a sequence-specific manner (Bell and Stillman, 1992). Concentrations of the helicase-loading proteins used were 10 nM Cdc6, 0.5 nM ORC, 10-15 nM Mcm2-7, and 10-20 nM Cdt1, with the exception of reactions in Figure 3 (see below). After helicase-loading proteins were perfused into the DNA-decorated chamber, 1-s duration images were acquired by alternate excitation with 532 and 633 nm lasers (laser powers were 0.7 mW and 0.35 mW respectively as measured incident to the micromirror). The average dead time between consecutive images was 0.2 s. The 2.4 s cycle was repeated 600 times in the course of an experiment (∼ 24 min).

### Single-molecule helicase loading assays mixtures of two labeled proteins

The helicase loading reactions described in Figs. 2 and 3 were performed with mixtures of N- and C-terminally labeled preparations of Mcm2-7 or ORC, respectively. In experiments with mixed Mcm2-7 preparations (Fig. 2), 7.5 nM Cdt1/Mcm2-7^2C-649^ and 7.5 nM Cdt1/Mcm2-7^4N-650^ were mixed and added to a reaction with 0.5 nM ORC and 10 nM Cdc6. In experiments with mixed ORC preparations, 0.6 nM ORC^1N-550^ was mixed with 0.25 nM ORC^5C-549^ because the bulk helicase loading activity of ORC^1N-550^ was lower than that of ORC^5C-549^. At the concentrations used, we observed equal frequencies of events where the first helicase is recruited by N- or C-terminally labeled ORC proteins (as determined by the FRET profiles and control experiments in which either ORC^5C-549^ or ORC^1N-550^ was the only donor protein), thus yielding the highest probability of potential “mixed” events. Standard concentrations of Cdt1/Mcm2-7^2C-649^ (15 nM) and Cdc6 (10 nM) were used.

### FRET Data Analysis

CoSMoS data sets were analyzed as described previously (Ticau et al., 2015), except that records of protein fluorescence were corrected for background fluorescence as described in (De Jesús-Kim et al., 2021). Records of acceptor-excited fluorescence in Figure 2 and Figure 3 were additionally corrected for spatial variations in local laser excitation as described previously (De Jesús-Kim et al., 2021). Donor-excited records (D_ex_, D_em_ and D_ex_, A_em_) used to calculate *E*_FRET_ were not subject to the corrections for local laser excitation in any instance.

By alternating between laser excitation wavelengths, we monitored the co-localization of both the donor fluorophore on ORC and acceptor fluorophore(s) on Mcm2-7 and/or Cdt1 with origin-DNA molecules. Double-hexamer formation events with no labeled ORC (donor) were excluded from data analysis. To determine the time of formation of the high-FRET state (*E*_FRET_ 0.7-0.8), we noted the earliest time the > 635 nm emission FRET signal rose above background noise level while a donor fluorophore was present (as determined by monitoring the signal in the > 635 nm field when only the 532 nm laser was turned on). Only spots where the arrival of the acceptor fluorophore could be confirmed within 4.8 s were considered genuine instances of high FRET. Because the events observed on each DNA molecule are independent measurements of a reaction, many replicates were combined in analysis of each experiment.

Apparent FRET efficiency was calculated using:

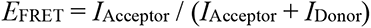

where *I*_Acceptor_ and *I*_Donor_ are the background-corrected acceptor and donor emission intensities, respectively, that are observed during donor excitation (De Jesús-Kim et al., 2021). Because the intensities are background-corrected, fluctuations in *I*_Acceptor_ and *I*_Donor_ sometimes result in *E*_FRET_ values below 0 or above 1. Rare *E*_FRET_ values greater than +2 or less than -2 were left out of the *E*_FRET_ histograms. Confidence bounds for kinetic data plotted as cumulative distributions were determined using the Greenwood formula as implemented in the Matlab ‘ecdf’ function.

### Determining ORC^5C-549^ labeling fraction

The fraction of ORC^5C-549^ molecules that were fluorescently labeled was determined as described previously (Ticau et al., 2015). Briefly, 10 μL (∼5.6 μg) of labeled ORC^5C-549^ was mixed with maleimide-DY-649P1 dissolved in anhydrous DMSO, in a 1:1 molar ratio at 4°C for 10 min. The reaction was terminated with 2 mM DTT. To ensure that we were monitoring fully assembled and functional ORC complexes, we added 1 nM of the double-labeled ORC to a single-molecule reaction chamber containing origin DNA and monitored ORC-DNA colocalization. The fraction of maleimide-DY-649P1-labeled ORC molecules that also contained DY-549P1 was determined and reported as the percent labeling by the DY-549P1.

### Determining ORC^5C-549^ photobleaching rate

To determine the photobleaching rate of ORC^5C-549^, single-molecule helicase loading reactions were performed with 0.5 nM ORC^5C-549^, 15 nM unlabeled Cdt1-Mcm2-7^5RA^, and 10 nM Cdc6 under two laser exposure conditions (Supplementary Figure 1h). Data for the experiments shown throughout the manuscript was acquired at a relative laser exposure of 1, by alternating between the two laser excitation wavelengths. For the ORC^5C-549^ photobleaching experiments, relative exposure of 1 was achieved by alternating acquisition of 1-s frames with excitation with the 532 nm laser and 1 s frames with no excitation (dead time between frames, 0.2s). The relative laser exposure was increased to 2.4 by continuously acquiring 1-s frames with 532 nm excitation and no dead time. ORC dwell times under the two laser exposures were each fit to single exponentials using maximum likelihood algorithms and bootstrap methods were used to determine uncertainty estimates, similar to what has been previously described (Crawford et al., 2013; Friedman and Gelles, 2012).

## Author Contributions

S.G. designed and conducted experiments with feedback from L.J.F., J.G., and S.P.B. S.G. analyzed data. S.G. and S.P.B. composed the paper with input from all authors, and S.P.B. and J.G. directed the project.

## Acknowledgements

We are grateful to members of the Bell laboratory for useful discussions. We thank Xiaoxue (Snow) Zhou and Hazal B. Kose for comments on the manuscript, Alexandra M. Pike and Paritosh Gangaramani for comments on the figures and Christian Ramsoomair for preparation of a subset of proteins described in this manuscript. This work was supported by NIH grants GM52339 (S.P.B.) and R01 GM81648 (J.G.). S.G. was supported in part by an NIH Pre-Doctoral Training Grant (T32 GM007287) and a MathWorks Science Fellowship. S.P.B. is an investigator with the Howard Hughes Medical Institute. We thank the Biopolymers core of the Koch Institute Swanson Biotechnology Center for technical support.

## Supplementary Figures

**Figure 1–figure supplement 1.**
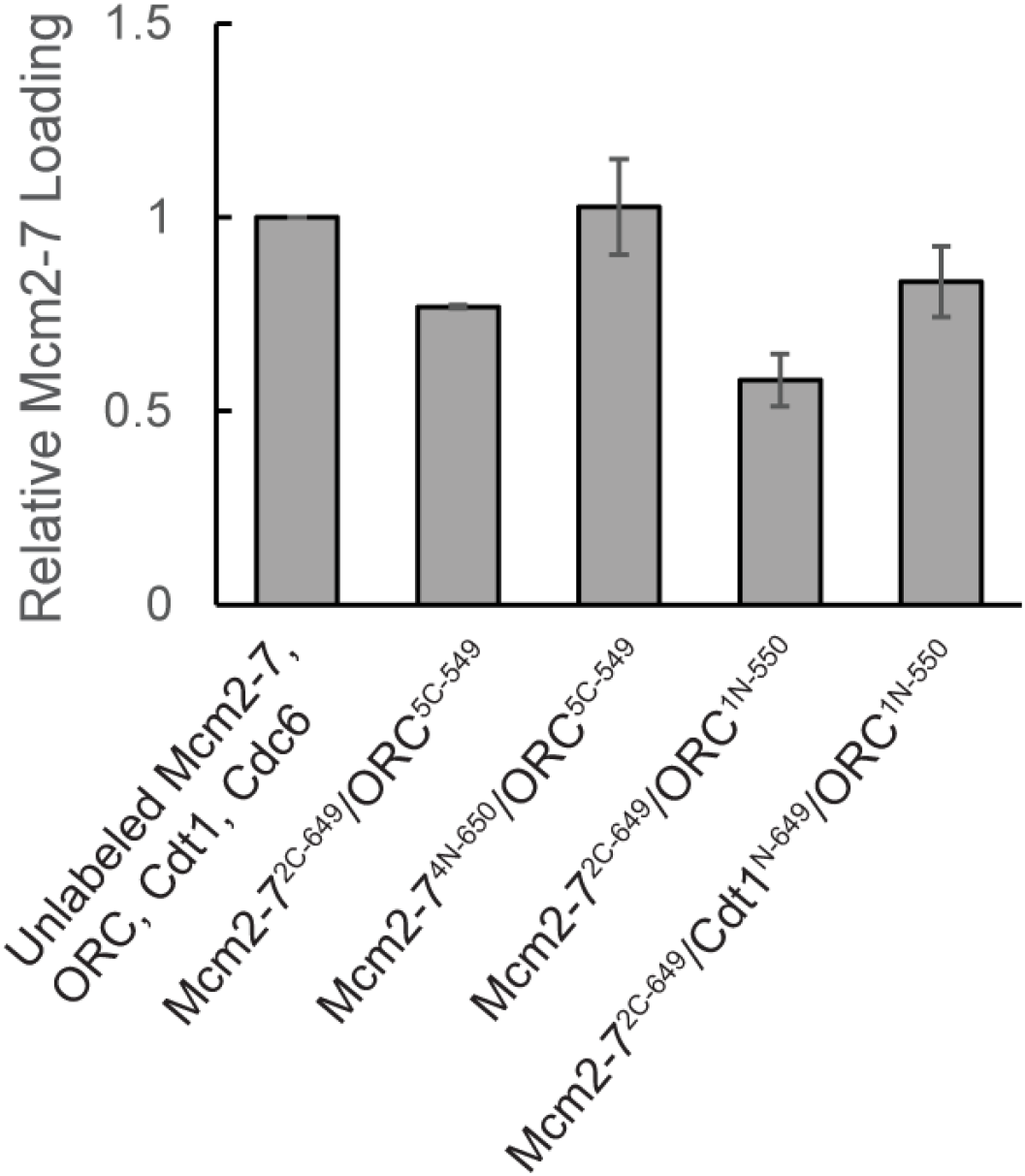
Fluorescently-labeled proteins function in ensemble helicase-loading assays. Bulk helicase-loading assays were used to measure the activity of the fluorescently-labeled proteins relative to their unlabeled counterparts. All samples were subjected to a high-salt wash, separated by SDS-PAGE, stained with Krypton protein stain, and quantified. Reactions were performed in duplicate and error bars indicate the standard deviation.

**Figure 1–figure supplement 2.**
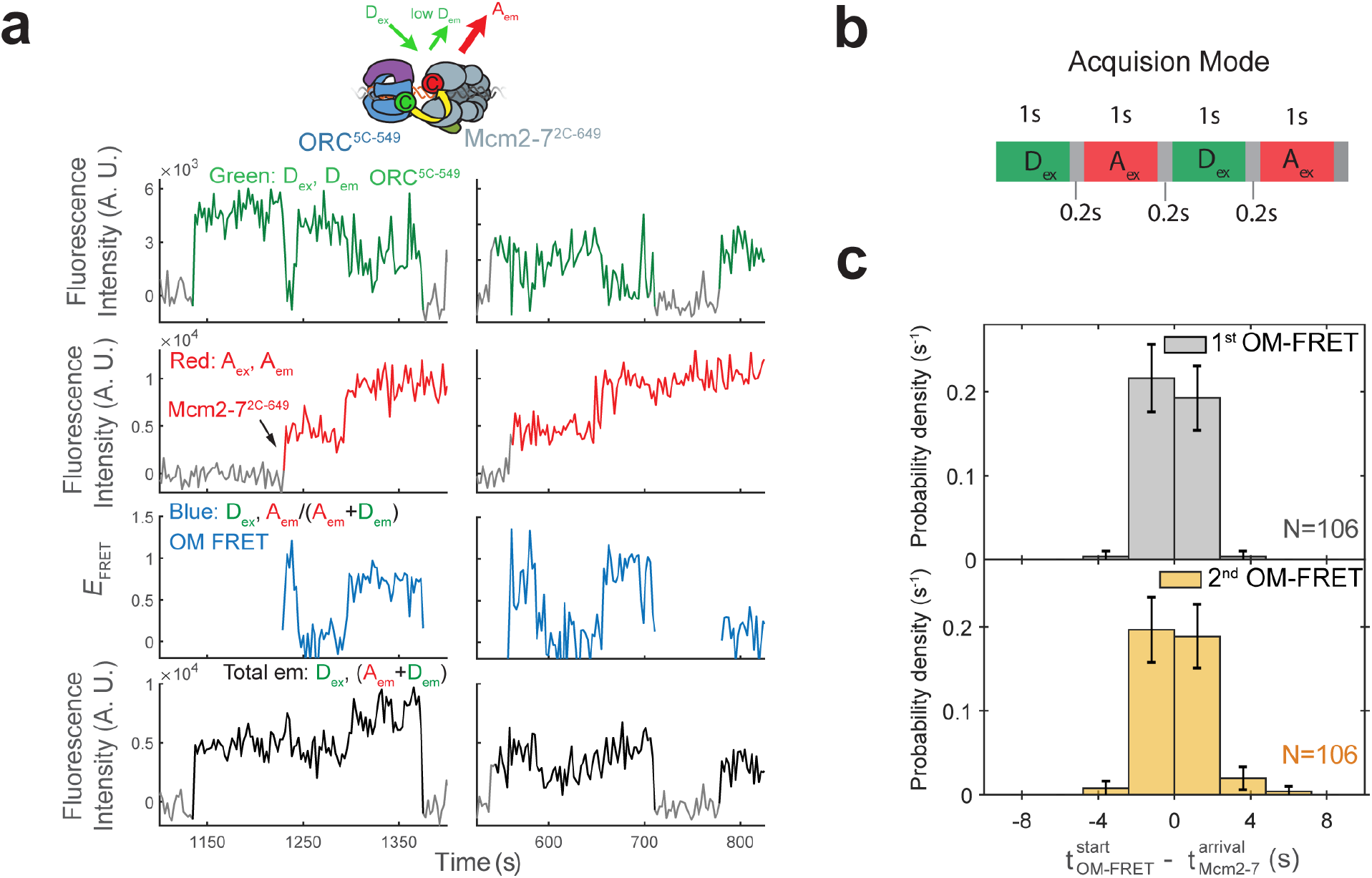
Each Mcm2-7 arrival is associated with an OM interaction. **(a)** Additional fluorescence intensity traces from the experiment in Figure 1c. **(b)** In our SM-FRET experiments, we acquired 1-s duration frames alternately exciting the donor and acceptor fluorophores with the 532 and 633 nm lasers, respectively. Gray boxes indicate dead time (average 0.2 s) between consecutive frames. **(c)** OM interactions begin when the corresponding Mcm2-7 associates with DNA. The time to OM-FRET after the corresponding Mcm2-7 arrival is plotted as a histogram. Top panel shows first OM interaction relative to first Mcm2-7 arrival and bottom panel shows second OM interaction relative to second Mcm2-7 arrival. OM-FRET start times within ± 2.4 s of Mcm2-7 arrival are simultaneous within the time resolution of our experiment. Negative values indicate instances where OM-FRET is detected in the donor-excited channel (D_ex_) before Mcm2-7 arrival is detected in the acceptor-excited channel (A_ex_); bin size is 2.4 s.

**Figure 1–figure supplement 3.**
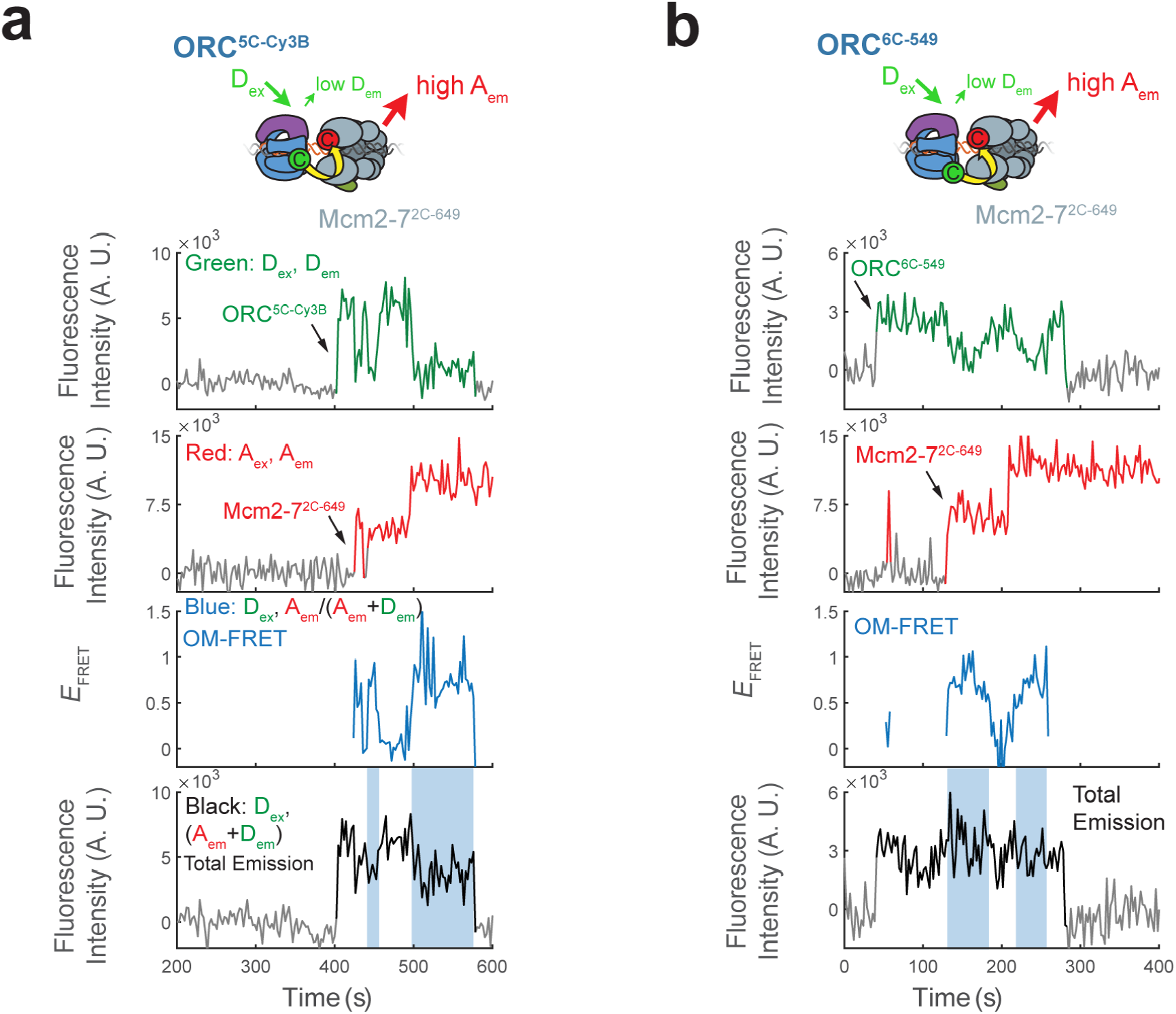
Enhancement of total emission upon donor excitation during OM interactions is due to protein-induced fluorescence enhancement (PIFE) exhibited by ORC^5C-549^. **(a)** The enhancement in total emission on donor excitation during periods of high *E*_FRET_ (Figure 1c, Black: Dex, Dem + Rem panel) is eliminated with an alternative dye (Hwang et al., 2011). The Cy3B fluorophore has a rigidified scaffold that does not exhibit PIFE (Hwang et al., 2011; Myong et al., 2009). Fluorescence intensity traces with ORC^5C-Cy3B^ show that when ORC is labeled at the same position as ORC^5C-549^ with the Cy3B fluorophore, total emission during OM-FRET is modestly reduced (compare with the total emission panels in Figure 1c and Figure 1– figure supplement 2a). Blue shaded areas indicate the times with OM-FRET during double-hexamer formation. **(b)** Enhanced total emission on donor excitation in experiments with ORC^5C-549^ (due to PIFE) is specific to the position of the label at the Orc5 C-terminus. When the position of the C-terminal label on ORC is changed from Orc5 to Orc6 but the same Dy549 dye is used we do not observe PIFE. Fluorescence intensity traces with ORC^6C-549^ show that the increase in total emission during OM-FRET is lost. Blue shaded areas indicate intervals with OM-FRET.

**Figure 1–figure supplement 4.**
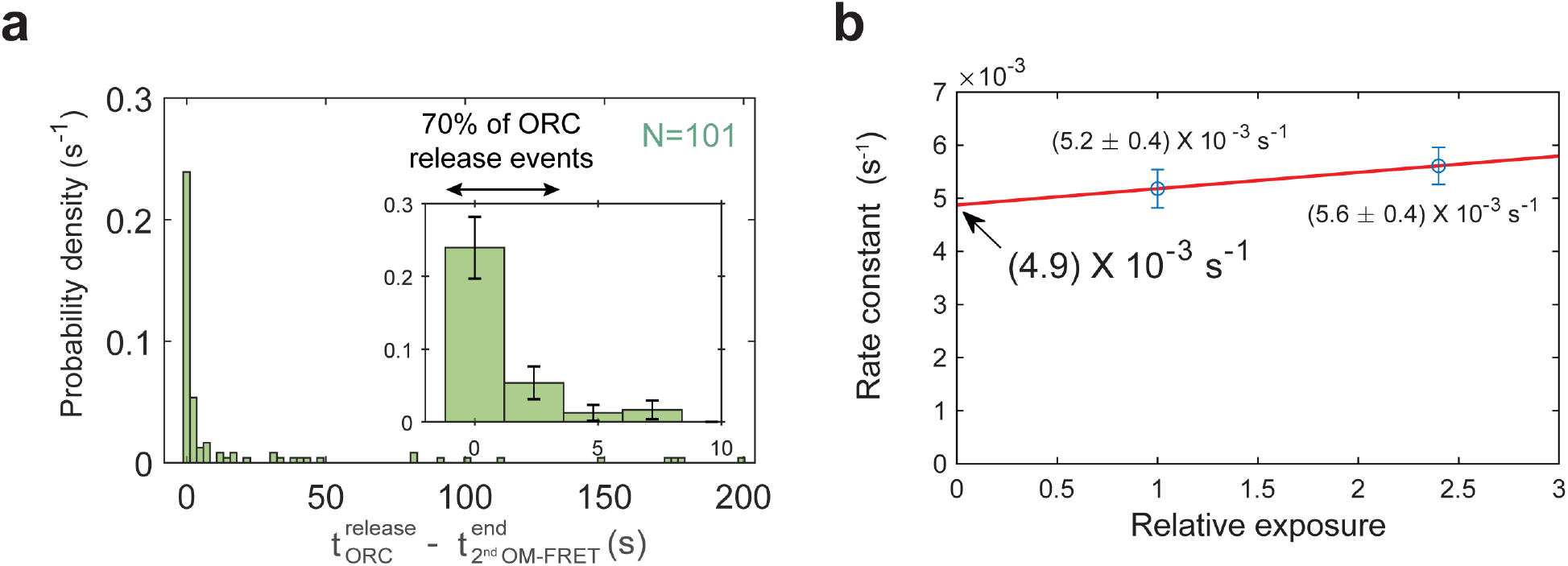
The end of the second OM interaction is concomitant with ORC departure in a majority of helicase loading events. **(a)** Histogram of time to ORC release after the second OM FRET ends. In 70% of the double-hexamer formation events, ORC releases within 3.6 s of the end of the second OM-FRET. Five long ORC release events are not shown (dwell times of 241 s, 266 s, 268 s, 382 s, and 417 s). Inset shows expanded x-axis and 74% of total events; bin size 2.4 s for both plots. **(b)** ORC^5C-549^ photobleaching rate measurement. The calculated dwell times for ORC^5C-549^ in a helicase-loading reaction were measured for different relative laser exposures and fit to single exponential functions. Data for the experiments shown throughout the manuscript were acquired at a relative laser exposure of 1. To measure the effect of photobleaching, the relative exposure was modulated by changing the exposure time to the laser. The relative exposure (x axis) is plotted against the off-rate constant (y axis ± SEM). ORC^5C-549^ photobleaching rate (under experimental conditions with relative exposure of 1 is (3.0 ± 3.6) x 10^-4^ s^-1^ which corresponds to a half-life of ∼2300 s. The mean dwell time of ORC^5C-549^ after second Mcm2-7 arrival is 88 ± 9 s. Because the photobleaching lifetime is much longer than the mean ORC dwell time (>26-fold longer), photobleaching does not appreciably contribute to the loss of ORC fluorescence observed in Figure 1c and Figure 1–figure supplement 2a.

**Figure 2–figure supplement 1.**
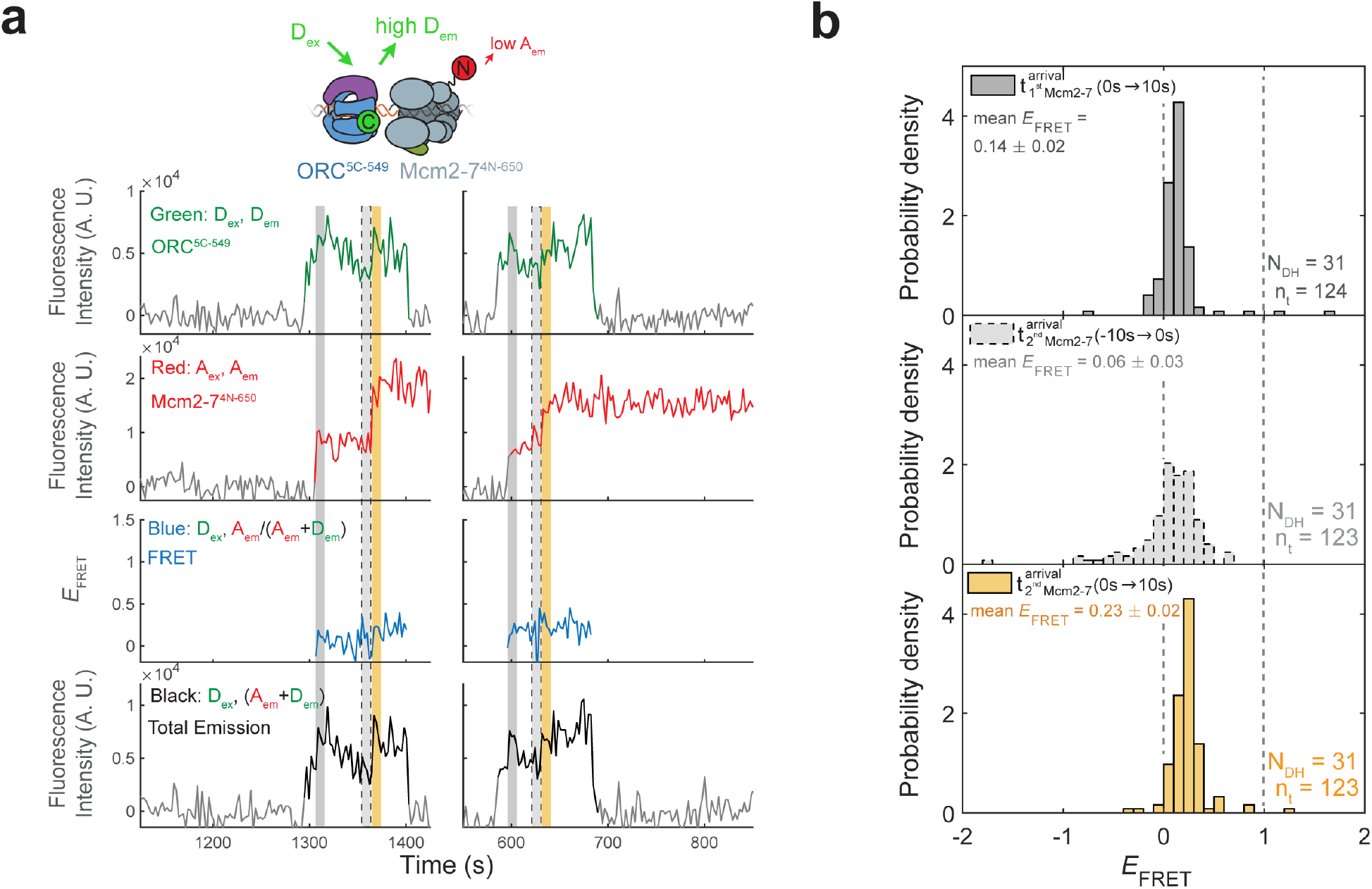
Mcm2-7^4N-650^ does not exhibit high *E*_FRET_ with ORC^5C-549^. **(a)** Fluorescence intensity traces from an experiment with ORC^5C-549^ (donor) and Mcm2-7^4N-650^ (acceptor). Note the low *E*_FRET_ values observed for this site of Mcm2-7 modification compared to those in Figure 1c. **(b)** Histogram plots of *E*_FRET_ values for 31 successful double-hexamer formation events with Mcm2-7^4N-650^ during the same three 10 s-time intervals described in Figure 1d: immediately after the first (top) or second Mcm2-7 (bottom) arrives or before the second Mcm2-7 arrives (middle). Examples of these intervals are indicated in (a) highlighted with gray, yellow, and gray dash, respectively. N_DH,_ number of double-hexamer formation events and n_t_, number of signal points.

**Figure 2–figure supplement 2.**
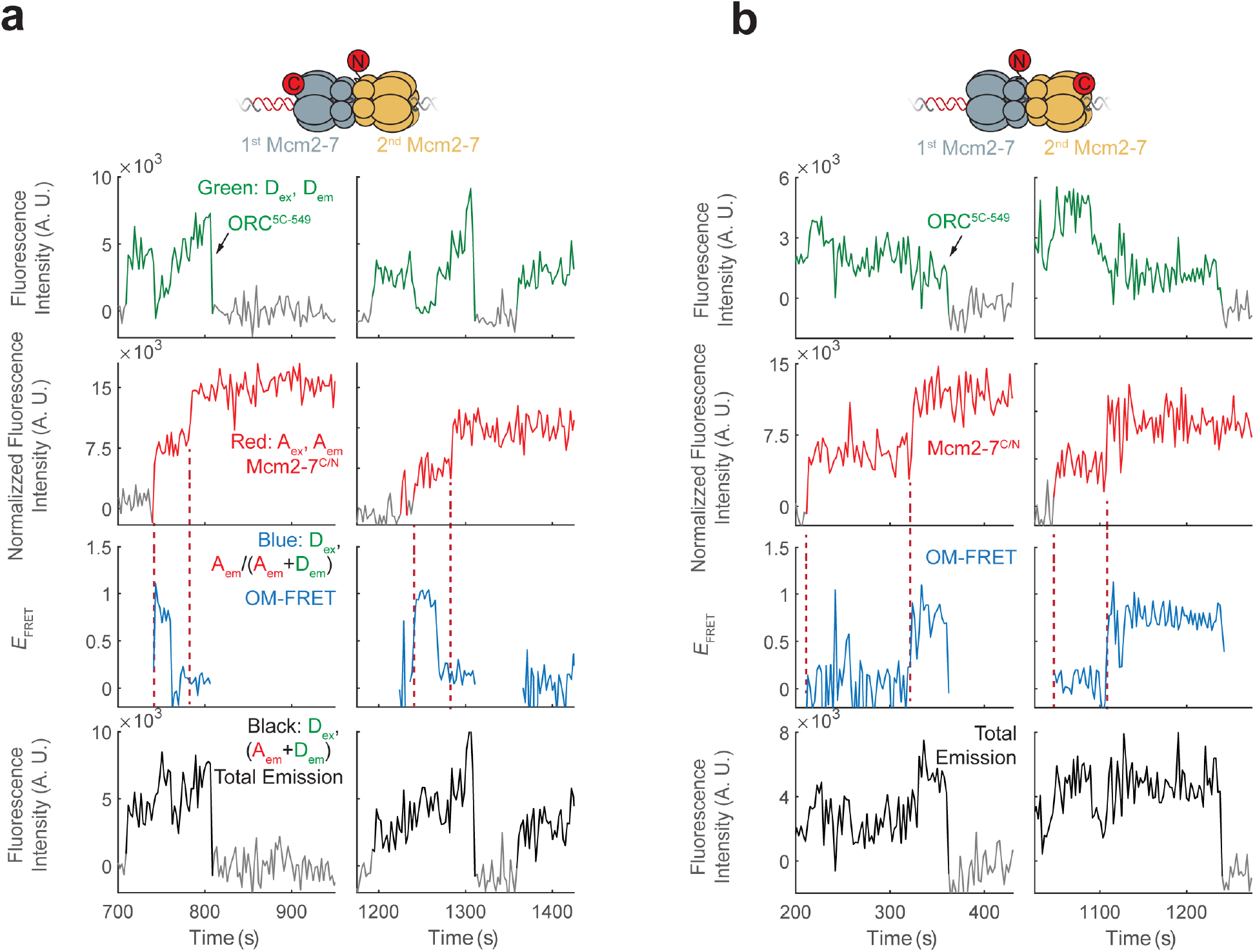
Additional fluorescence intensity traces for the mixed double hexamer populations with single OM FRET peaks observed in. Figure 2c**. (a)** Two examples of CN double hexamers where only the first Mcm2-7 arrival is accompanied by an OM interaction. **(b)** Two examples of NC double hexamers in which only the second Mcm2-7 arrival is accompanied by an OM interaction.

**Figure 3–figure supplement 1.**
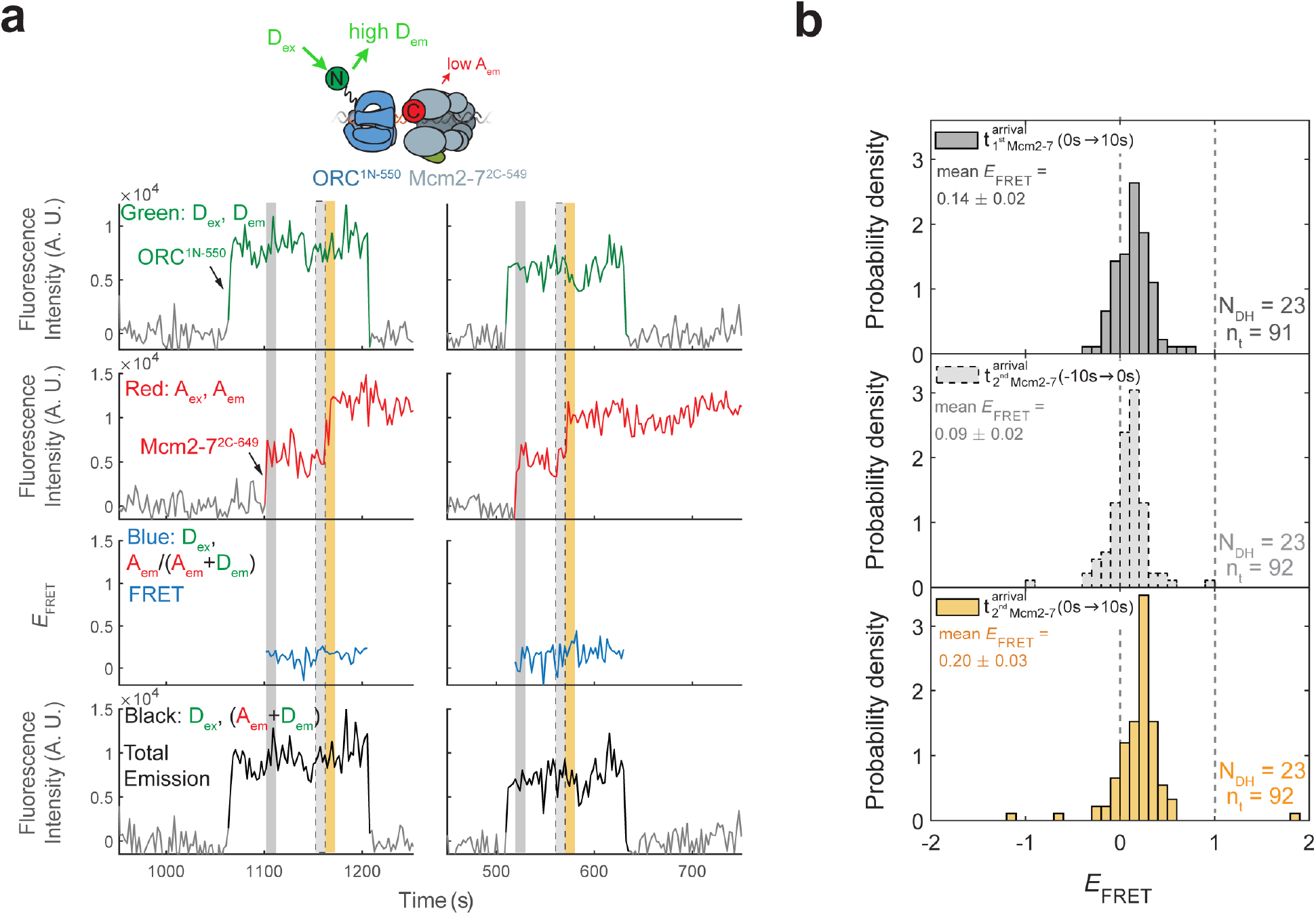
**ORC^1N-550^ does not exhibit high *E*_FRET_ with Mcm2-7^2C-649^. (a)**. Fluorescence intensity traces from an experiment with ORC^1N-550^ which is labeled at the N-terminus of Orc1 with a donor fluorophore and Mcm2-7^2C-649^ (acceptor fluorophore). Note the low *E*_FRET_ values observed for this site of ORC modification compared to those in Figure 1c. **(b)** Histogram plots of *E*_FRET_ values for 23 successful double-hexamer formation events with ORC^1-550^ during the same three 10s-time intervals in Figure 1d: immediately after the first (top) or second Mcm2-7 (bottom) arrives or before the second Mcm2-7 arrives (middle). Examples of these intervals are indicated in (a) highlighted with gray, yellow, and gray dash, respectively. N_DH,_ number of double-hexamer formation events and n_t_, number of signal points.

**Figure 4–figure supplement 1.**
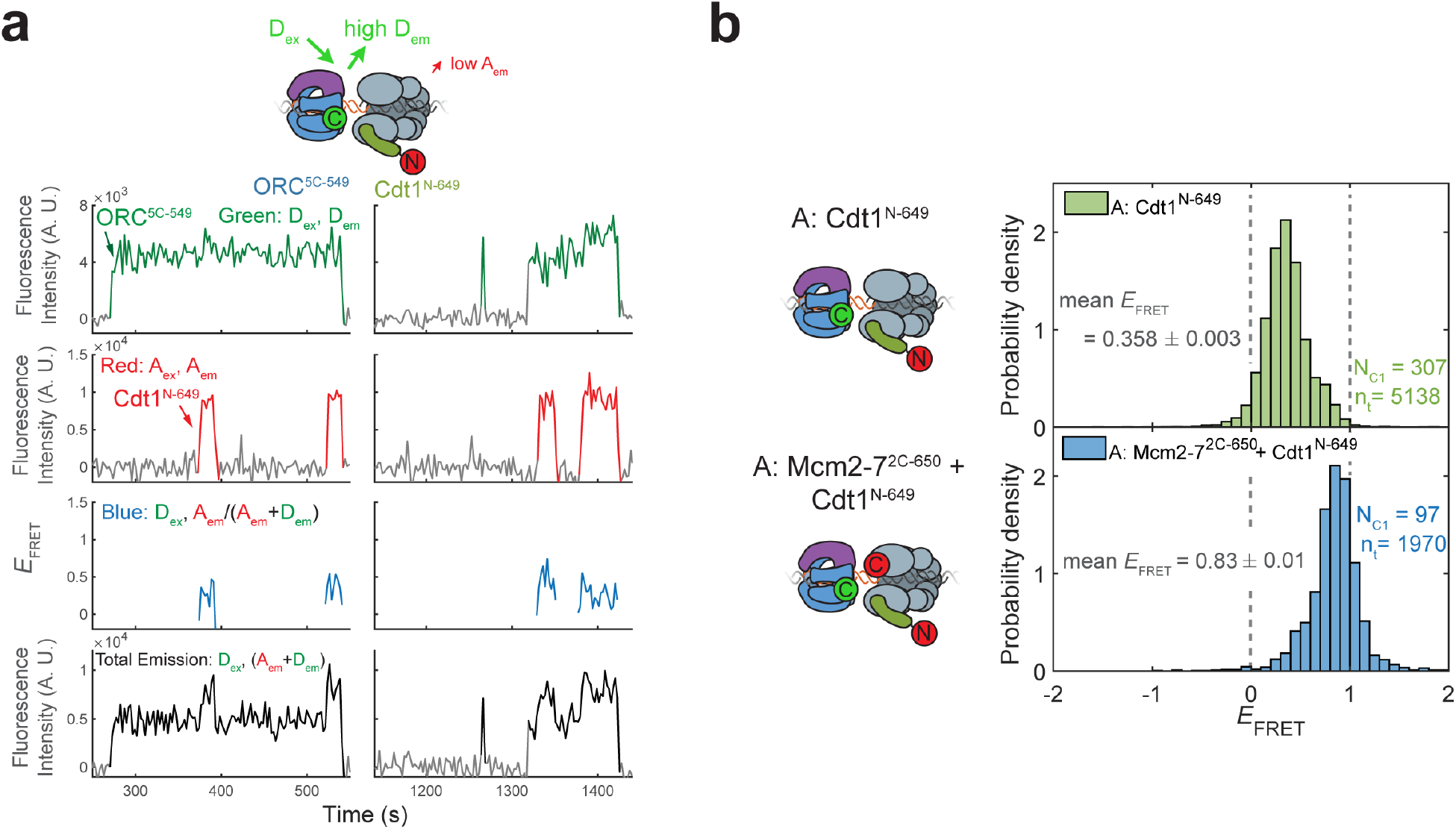
Cdt1^N-649^ does not exhibit high *E*_FRET_ with ORC^5C-549^. **(a)** Fluorescence intensity traces for an experiment with ORC^5C-549^ (donor), Cdt1^N-649^ (acceptor), and unlabeled Mcm2-7. Although FRET is observed when Cdt1 is present, the values are consistently lower than that observed between ORC^5C-549^ and Mcm2-7^2C-649^. **(b)** Histograms of *E*_FRET_ values with ORC^5C-549^, Cdt1^N-649^, and unlabeled Mcm2-7 (top) compared with reactions in which Mcm2-7 is also labeled with an acceptor fluorophore at the Mcm2 C-terminus (Mcm2-7^2C-650^, bottom). The two dashed lines indicate *E*_FRET_ values of 0 and 1. N_C1,_ number of Cdt1 binding events and n_t_, number of signal points.

**Figure 4–figure supplement 2.**
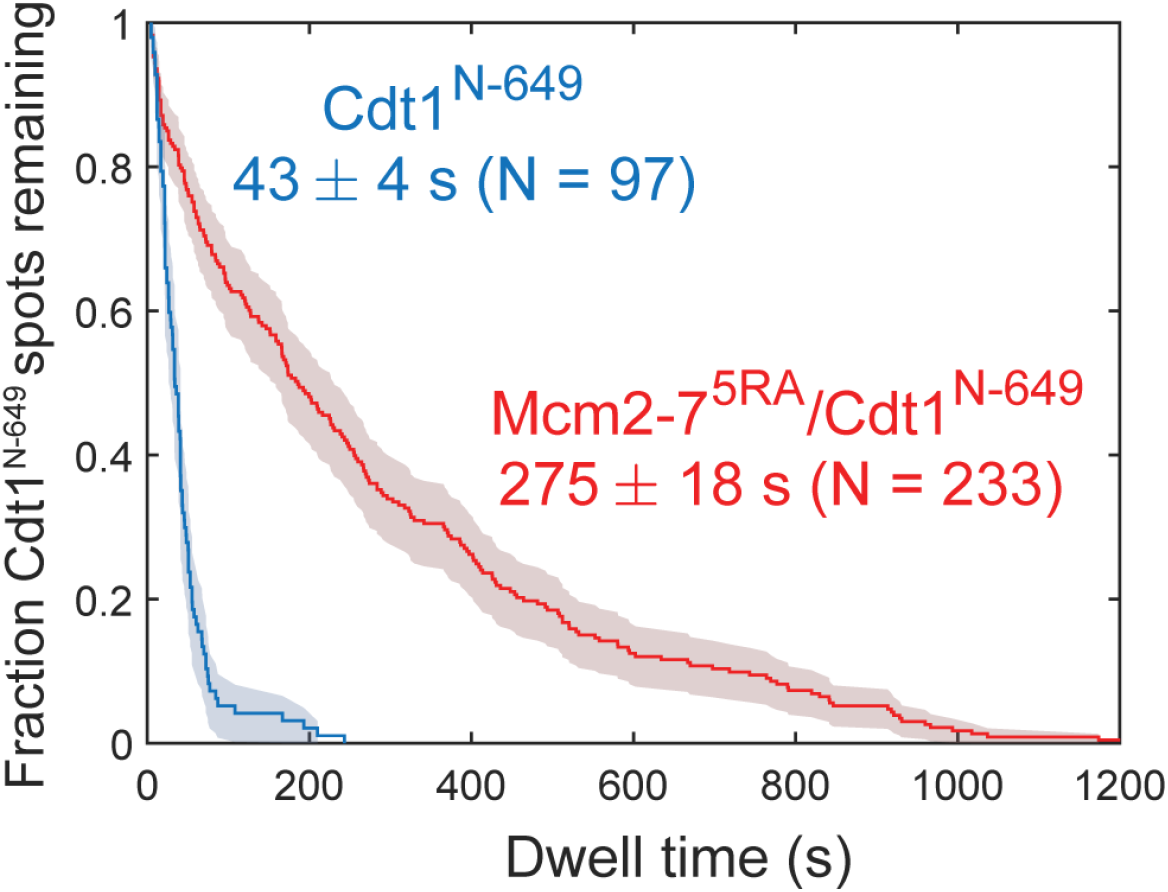
Disappearance of Cdt1^N-649^ fluorescence in the experiment in Figure 4b, 4c cannot be explained by photobleaching. Comparison between the distribution of Cdt1^N-649^ dwell times after the first Mcm2-7 arrival from the experiment in Figure 4c (wild-type) and an otherwise identical experiment with the Mcm2-7^5RA^ mutant that does not release Cdt1. The mean Cdt1 dwell time seen with the mutant sets a lower limit on the photobleaching lifetime. The mean Cdt1 dwell time in the wild-type experiment is >6-fold shorter, demonstrating that photobleaching does not appreciably contribute to the acceptor fluorescence decrease in the wild-type experiment.

**Figure 4–figure supplement 3.**
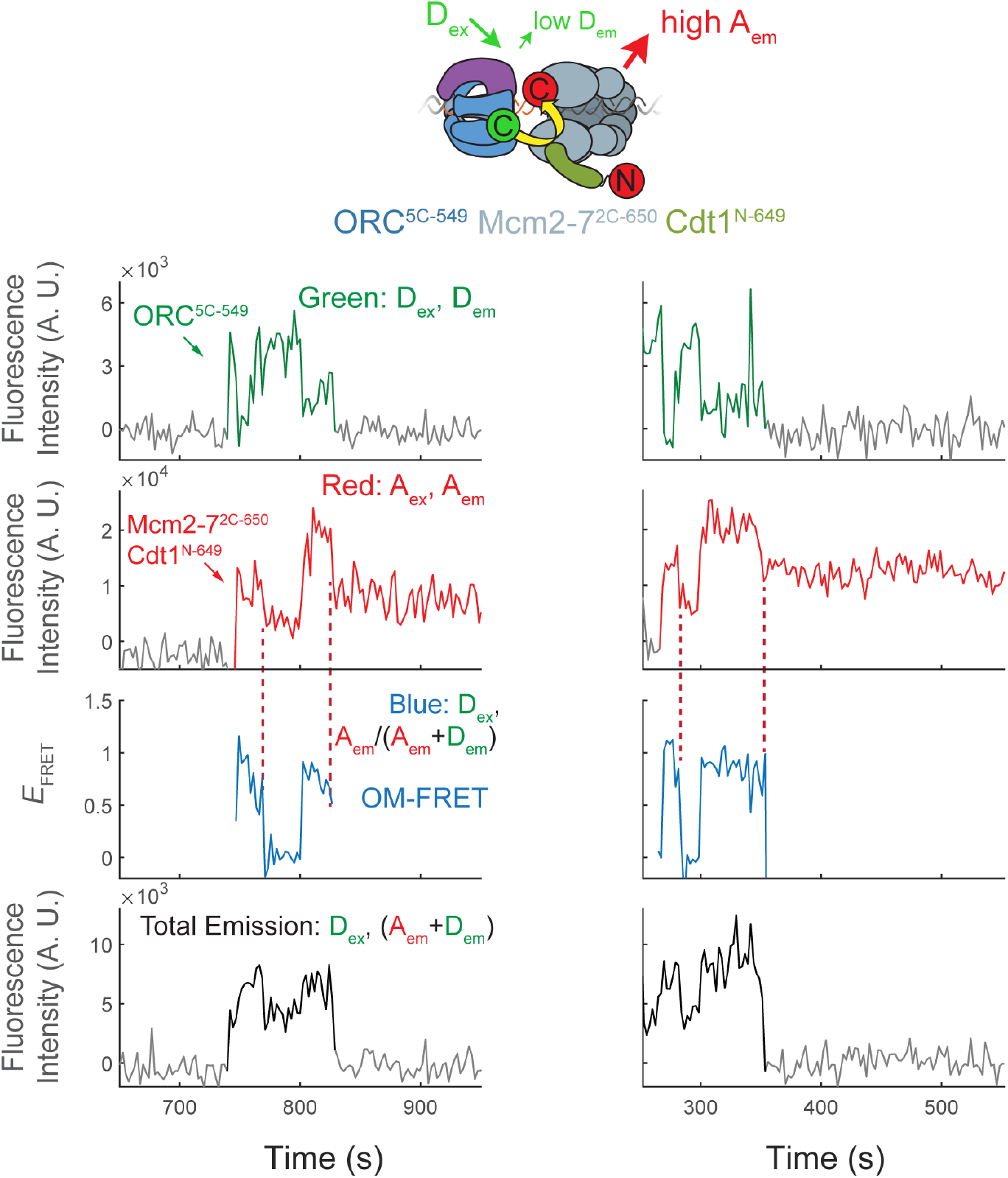
Representative traces show Cdt1 release corresponds with the end of OM interactions on individual DNAs. Additional fluorescence intensity traces from the experiments shown in Figure 4b and 4c.

**Figure 4–figure supplement 4.**
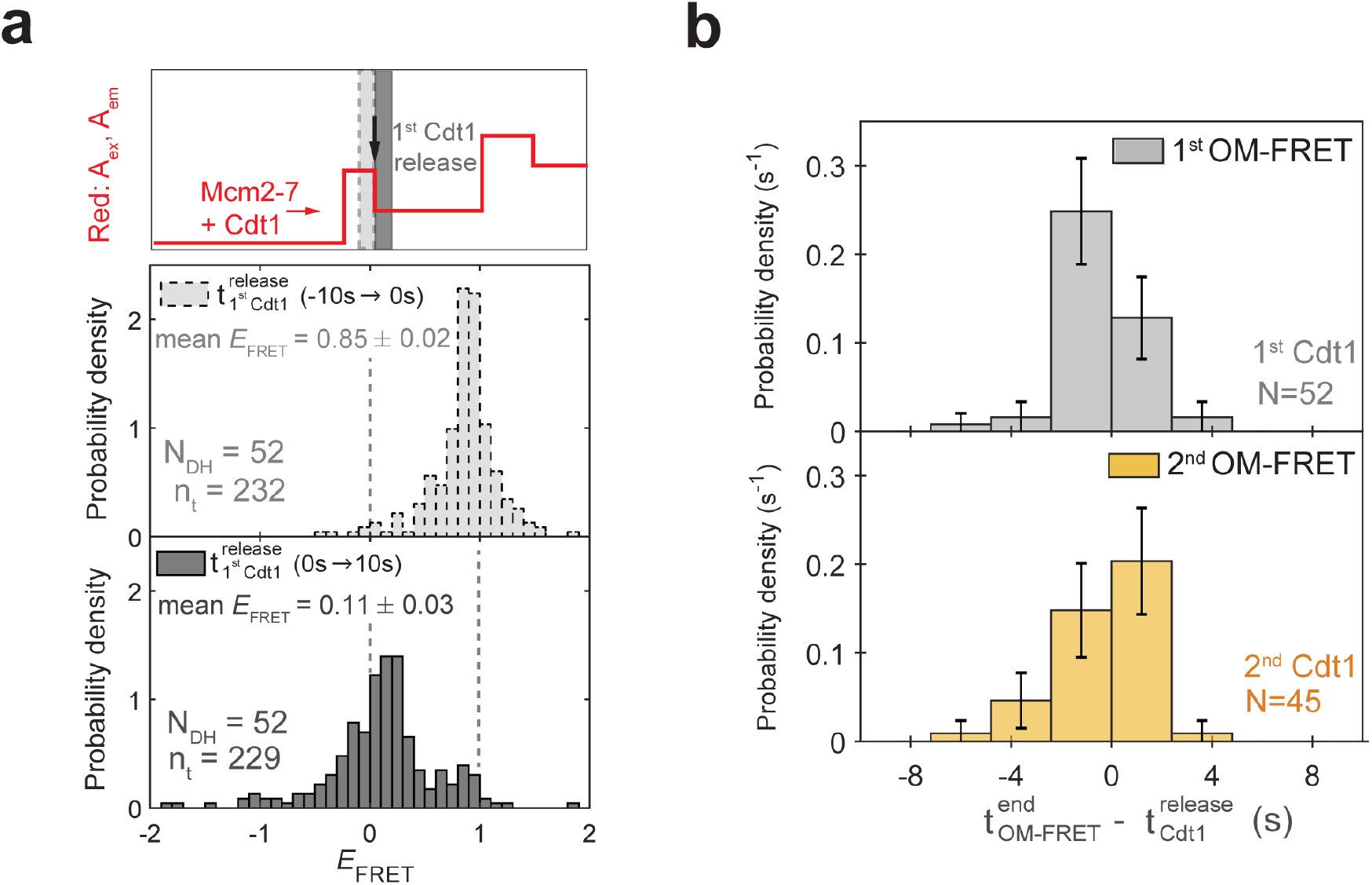
Cdt1 release is concomitant with loss of the high OM *E*_FRET_. **(a)** Distributions of *E*_FRET_ values from the experiment in Figures 4b and 4c. Top: Schematic of predicted Red: A_ex_, A_em_ signal for two Mcm2-7^2C-650^/Cdt1^N-649^ association events followed by Cdt1^N-649^ release that result in double-hexamer formation. The black arrow indicates the first Cdt1^N-649^ release, which results in loss of half the acceptor fluorescence. Middle, bottom: Histograms of *E*_FRET_ values for the two 10s-time intervals immediately before (middle) or after (bottom) 52 first Cdt1-release events. The two dashed lines indicate *E*_FRET_ values of 0 and 1. N_DH,_ number of double-hexamer formation events and n_t_, number of signal points. **(b)** Most first and second Cdt1 release events end at the same time as the corresponding OM interactions. The time of OM-FRET end after the corresponding Cdt1 release (±SE) is plotted as a histogram. Top panel shows first OM interaction relative to first Cdt1 release and bottom panel shows second OM interaction relative to second Cdt1 release.

**Figure 5–figure supplement 1.**
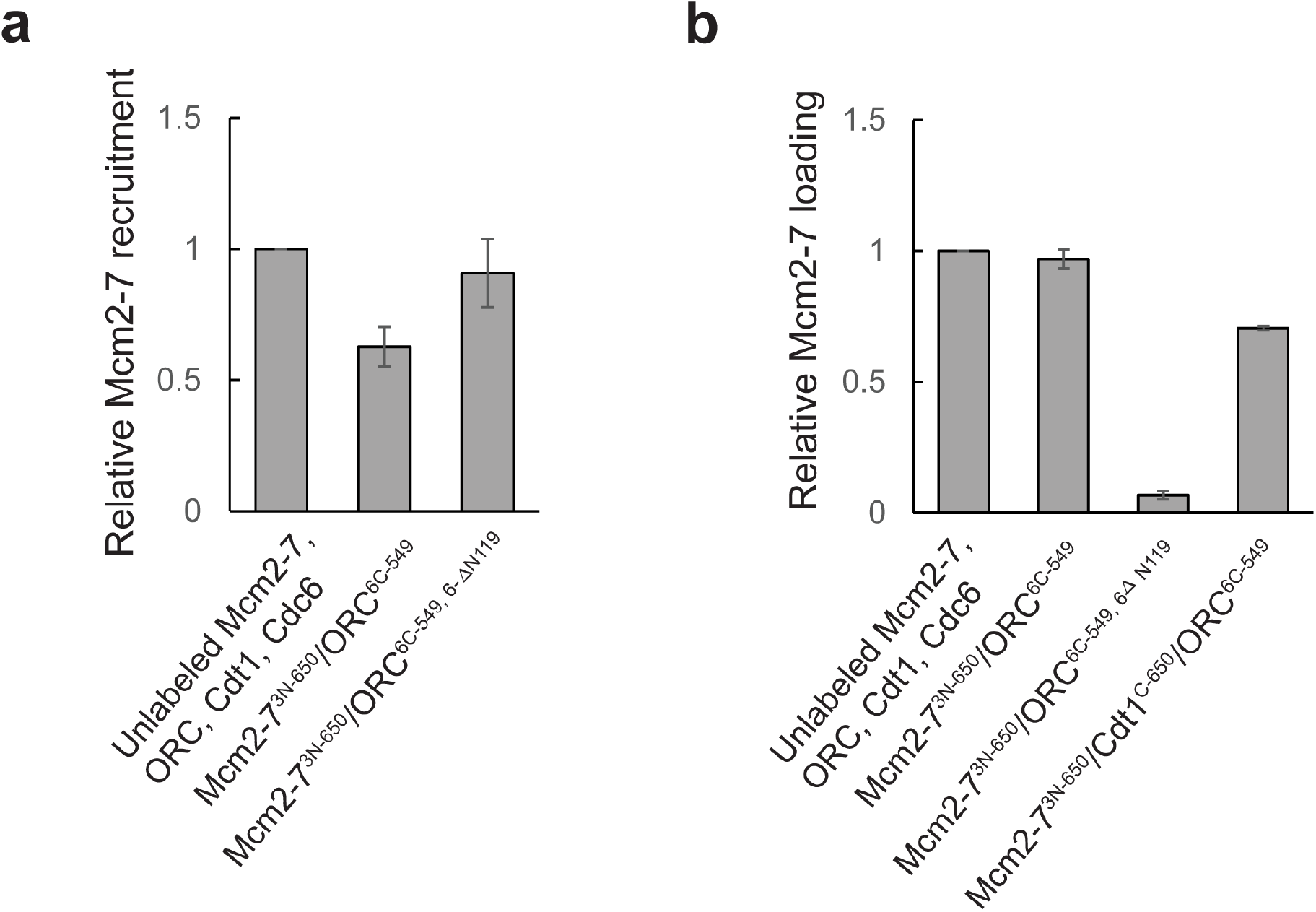
Fluorescent labels used for detecting the MO interaction function in ensemble helicase-loading assays. Activity of fluorescently labeled proteins relative to the unlabeled proteins in **(a)** ensemble helicase recruitment and **(b)** ensemble helicase loading assays. Helicase recruitment assay was performed in the same manner as the helicase-loading assay except ATP was replaced with the slowly-hydrolyzed ATP-analog ATPγS, and samples were not subjected to a high-salt wash. All samples were separated by SDS-PAGE, stained with Krypton protein stain, and quantified. Reactions were performed in duplicate and error bars indicate the standard deviation. Note that, as expected, ORC^6C-549,6-ΔN119^ has a helicase loading defect, but is not defective for helicase recruitment (Miller et al., 2019).

**Figure 5–figure supplement 2.**
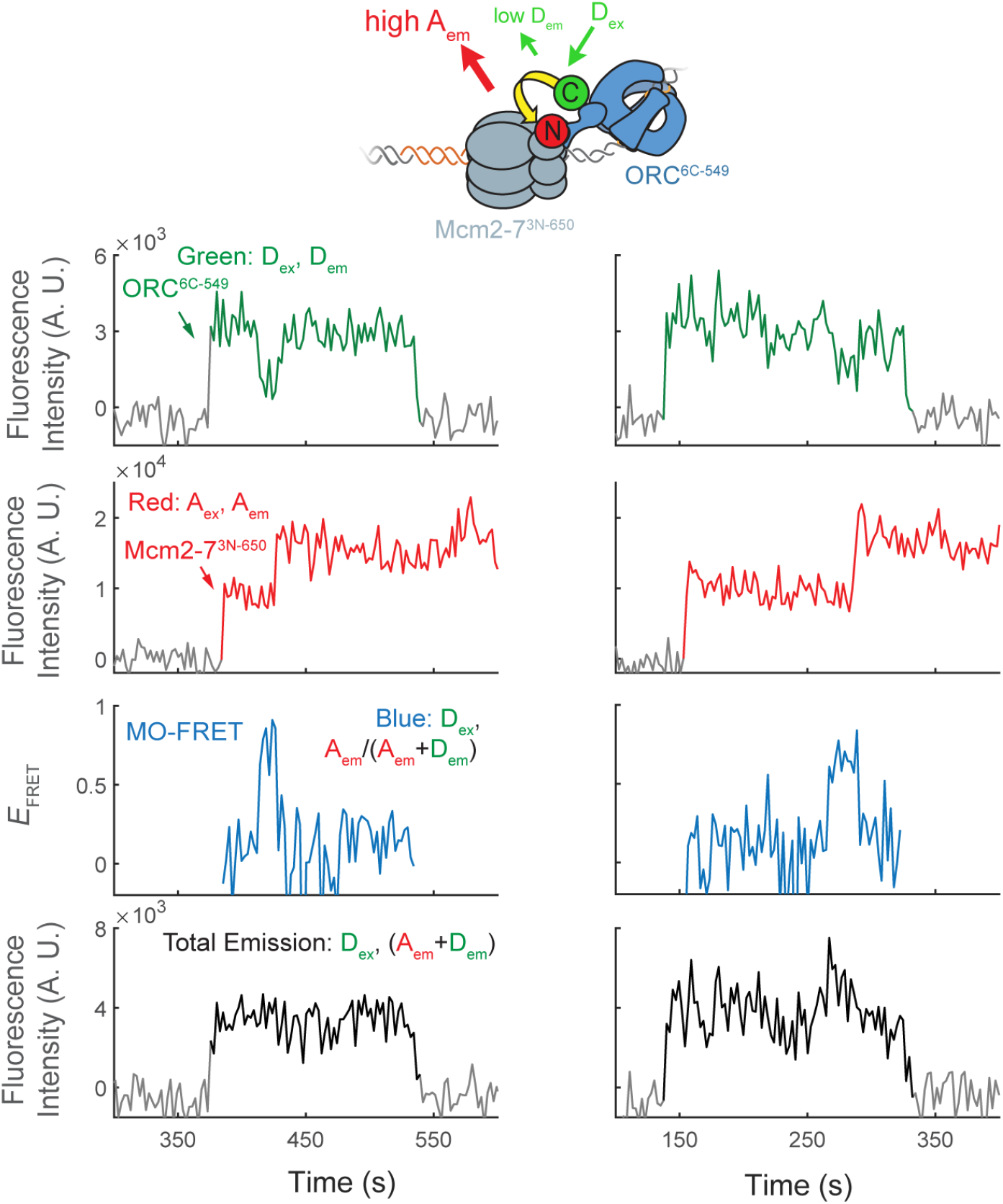
Representative traces showing MO interactions during helicase loading. Additional fluorescence intensity traces for the experiment in Figure 5c.

**Figure 5–figure supplement 3.**
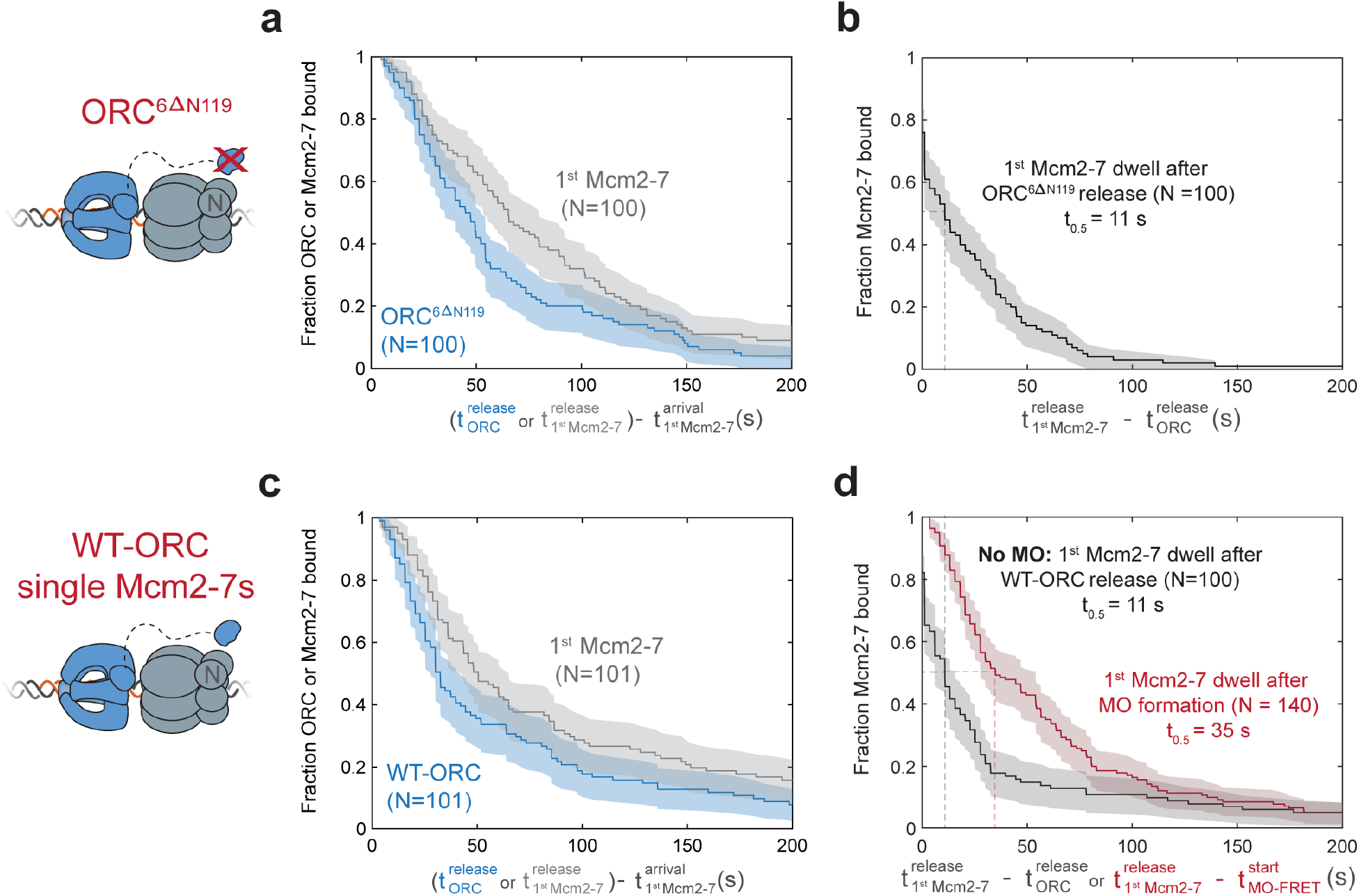
First Mcm2-7 helicases that do not form MO complexes are rapidly released. **(a) and (b)** In the context of the ORC^6C-549,6ΔN119^ mutant, the first Mcm2-7 dissociates rapidly after ORC is released. **(a)** Dwell times of ORC^6C-549,6ΔN119^ (blue; ORC^6ΔN119^) and Mcm2-7^3N-650^ (gray) after first Mcm2-7 arrival are plotted as survival functions. **(b)** Dwell times of Mcm2-7^3N-650^ after ORC^6C-549,6ΔN119^ is released from the DNA are plotted as a survival function. **(c) and (d)** In the context of wild-type ORC^6C-649^, first Mcm2-7 helicases that do not form an MO complex dissociate rapidly after ORC is released. **(c)** Dwell times of ORC^6C-549^ (blue; WT-ORC) and Mcm2-7^3N-650^ (gray) after first Mcm2-7 arrival for loading events that do not form MO interactions (from experiment in Figure 5c) are plotted as survival functions. **(d)** Dwell times of the Mcm2-7^3N-650^ molecules that do (red) or do not (black) form MO interactions are plotted. First Mcm2-7 dwell times that do not form MO interactions are plotted relative to release of ORC from the DNA (black). First Mcm2-7 dwell times in cases where MO interactions are formed are plotted relative to time of MO-FRET initiation (red). In events where a first Mcm2-7 goes on to recruit a second Mcm2-7, time of second Mcm2-7 arrival is used as a lower bound on the time the first Mcm2-7 is released. Shaded areas represent the 95% confidence intervals for each curve and t_0.5_ denotes the time at which half the Mcm2-7 molecules have been released. Sample size was limited to the first 100/410 Mcm2-7 associations in ORC^6C-549,6ΔN119^ data (a, b) and to the first 101/509 Mcm2-7 associations in wild-type ORC^6C-649^ data (c, d).

**Figure 6–figure supplement 1.**
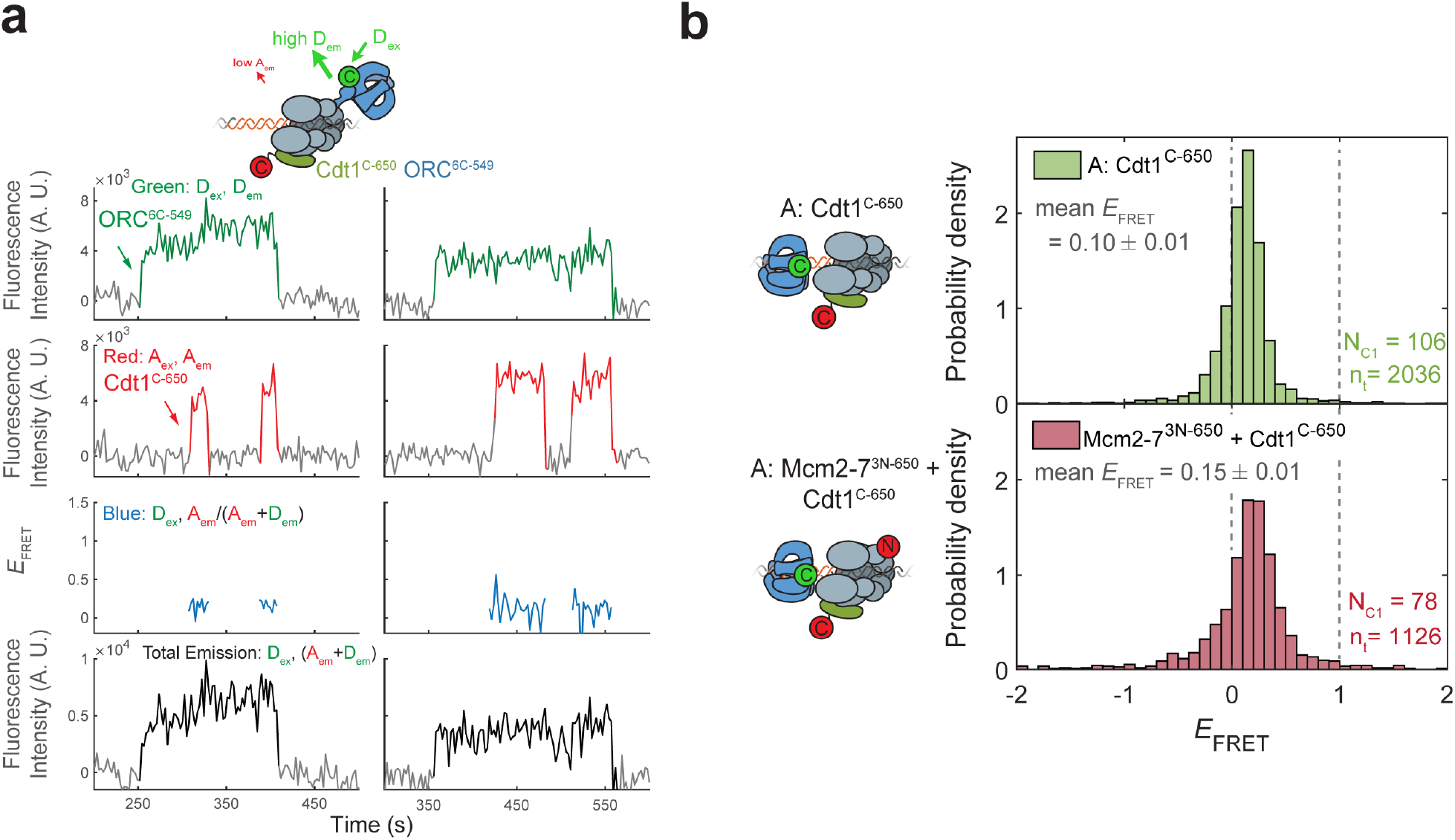
Cdt1^C-650^ does not exhibit high *E*_FRET_ with ORC^6C-549^. **(a)** Fluorescence intensity traces for an experiment with ORC^6C-549^ (donor), Cdt1^C-650^ (acceptor), and unlabeled Mcm2-7. **(b)** Histogram plots of *E*_FRET_ values for the duration that Cdt1 is associated in experiments with ORC^6C-549^. Top panel: Cdt1^C-650^ and unlabeled Mcm2-7 and bottom panel: Cdt1^C-650^ and Mcm2-7^3N-650^ are both labeled. Note that MO-*E*_FRET_ is low when Cdt1 is associated because MO-FRET is initiated after Cdt1 release (compare with Figure 6–figure supplement 3b). N_C1,_ number of Cdt1 binding events and n_t_, number of signal points.

**Figure 6–figure supplement 2.**
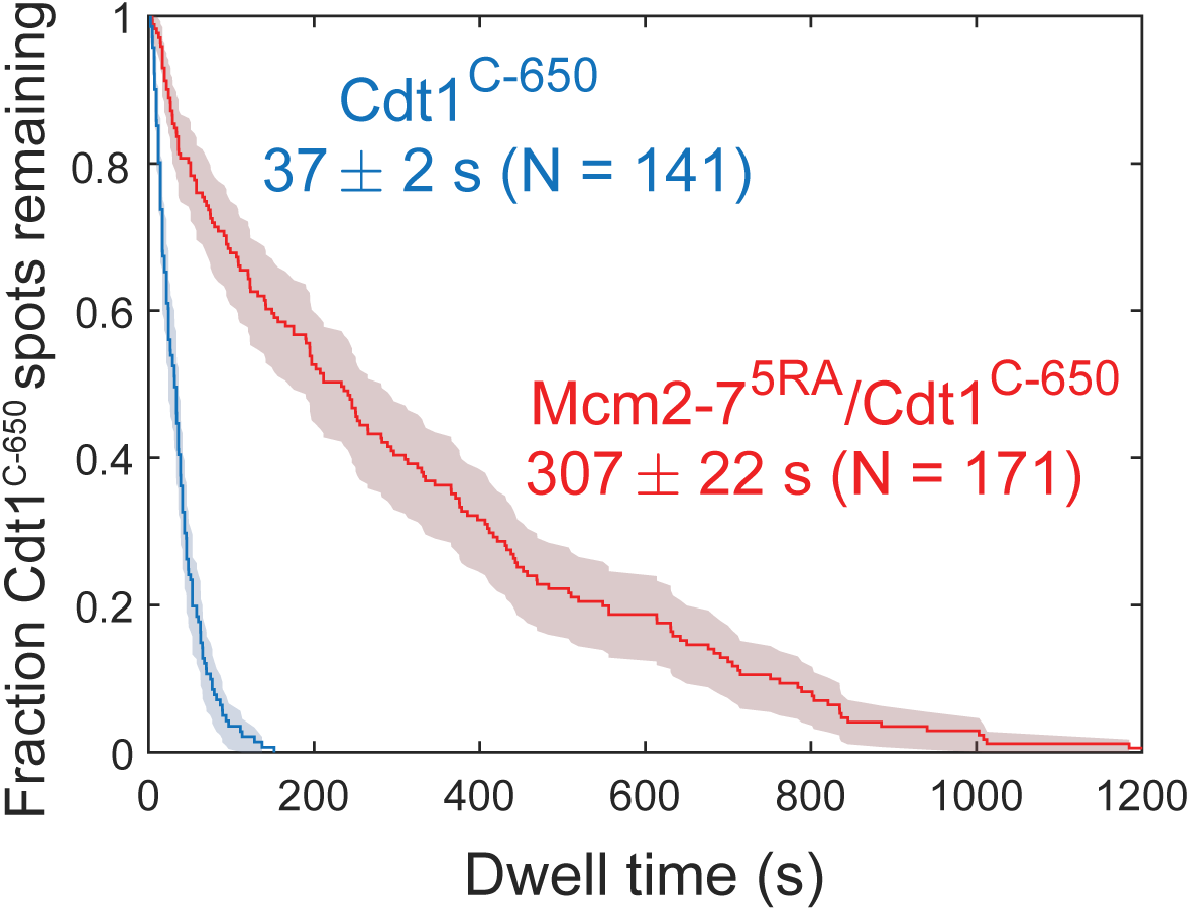
Disappearance of Cdt1^C-650^ fluorescence in the experiment in Figure 6a and 6b cannot be explained by photobleaching. Comparison between the distribution of Cdt1^C-650^ dwell times after the first Mcm2-7 arrival from the experiment in Figure 6b (wild-type) and an otherwise identical experiment with the Mcm2-7^5RA^ mutant that does not release Cdt1. The mean Cdt1 dwell time seen with the mutant sets a lower limit on the photobleaching lifetime. The mean Cdt1 dwell time in the wild-type experiment is >8-fold lower, demonstrating that photobleaching does not appreciably contribute to the acceptor fluorescence decrease in the wild-type experiment.

**Figure 6–figure supplement 3.**
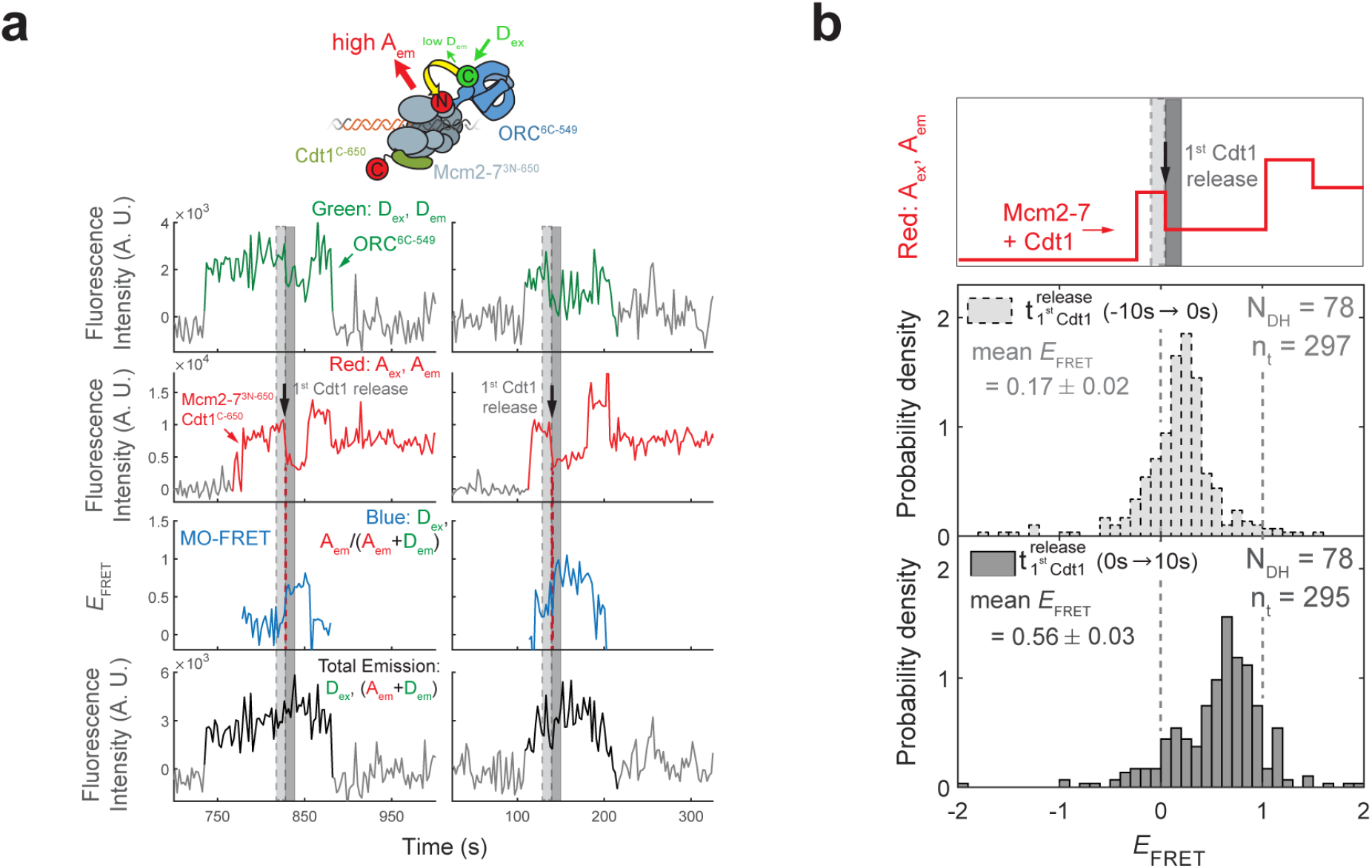
The high MO *E*_FRET_ state is established rapidly after the first Cdt1 is released. **(a)** Additional examples of fluorescence intensity traces for the experiment in Fig. 6c. **(b)** *E*_FRET_ values in 10s intervals on either side of first Cdt1 release in the experiment shown in Figure 6a. The example trace (Red: A_ex_, A_em_ panel on top) shows two Mcm2-7^2C-650^/Cdt1^C-650^ association events followed by Cdt1^C-650^ release that result in double-hexamer formation. The black arrow indicates the first Cdt1^C-650^ release, which results in loss of half the acceptor fluorescence. Histogram plots of *E*_FRET_ values for two 10s-time intervals prior to (middle) or after (bottom) 78 first Cdt1 release events. Examples of these intervals are indicated in the top panel highlighted with gray-dash and gray respectively. The two dashed lines indicate *E*_FRET_ values of 0 and 1. N_DH,_ number of double-hexamer formation events and n_t_, number of signal points.

**Figure 6–figure supplement 4.**
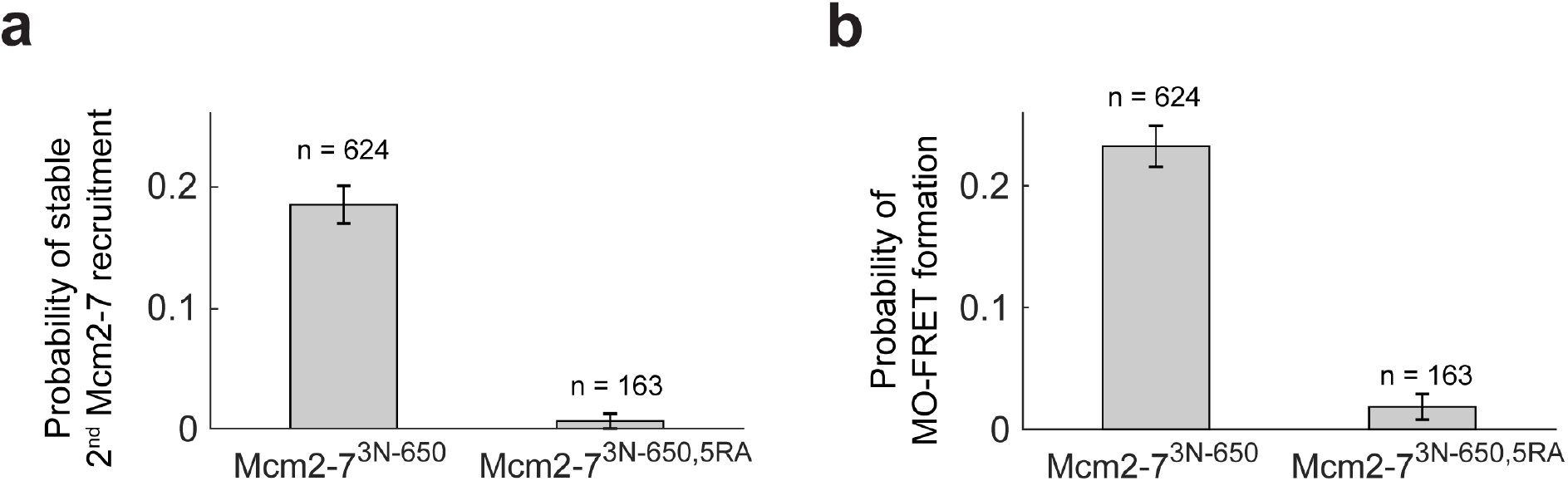
The first Cdt1 release anticipates formation of the MO interaction. **(a)** Preventing first Cdt1 release interferes with second Mcm2-7 recruitment. The fraction (± SE) of n first Mcm2-7^3N-650^ binding events that resulted in stable recruitment of a second Mcm2-7^3N-650^ is plotted for WT Mcm2-7 and Mcm2-7^5RA^ proteins. **(b)** Prevention of first Cdt1 release interferes with the establishment of the MO interaction. The fraction (± SE) of n first Mcm2-7^3N-650^ binding events that resulted in MO-FRET formation is plotted for WT-Mcm2-7 and Mcm2-7^5RA^ proteins.

**Figure 7–figure supplement 1.**
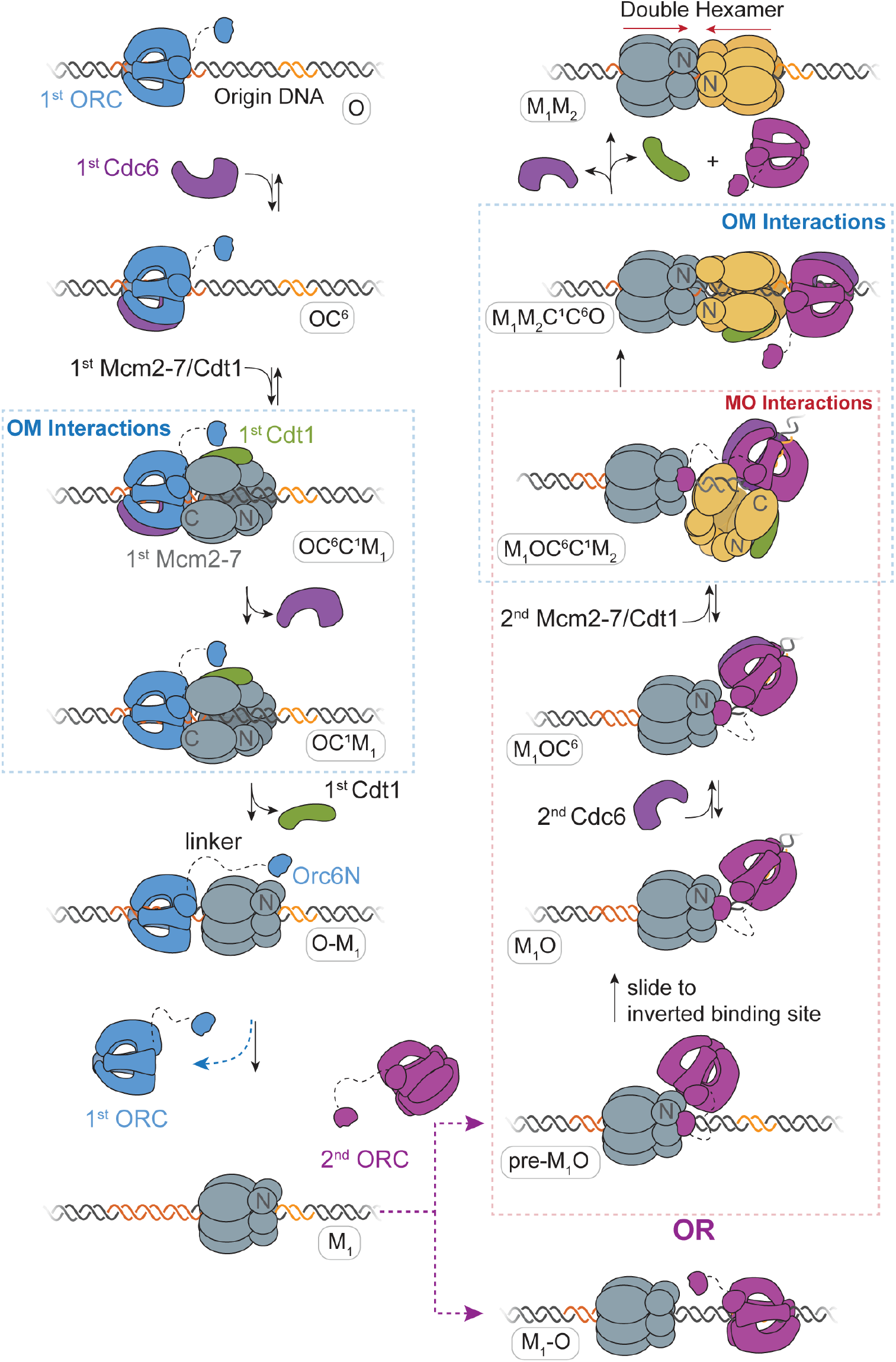
A model for helicase loading mediated by two separate ORC molecules. Designations in ovals are unique names for each intermediate using the abbreviations O, ORC; C^6^, Cdc6; C^1^, Cdt1; M_1_, 1^st^ Mcm2-7; and M_2_, 2^nd^ Mcm2-7. Blue and purple coloring distinguishes the ORC molecules that guide the first and second helicase recruitment respectively. Note that O_1_-M and M_1_-O are states where the ORC and Mcm2-7 are DNA-bound but not interacting.

**Supplementary Table 1.** Yeast Strains Used in this Study.

| **Strain** | **Additional information** | **Genotype** | **Source** |
| --- | --- | --- | --- |
| yMH09 | Base Strain | *MATa ade2-1 trp1-1 leu2-3,112 his3-11,15 ura3-1 can1-100 bar1::HisG lys2::HisG pep4Δ::unmarked* | Bell Lab |
| yRH101 | Base Strain | *MATa ade2-1 trp1-1 leu2-3,112 his3-11,15 ura3-1 can1-100 bar1::HisG lys2::HisG pep4Δ::KanMX* | Bell Lab |
| yST210 | Mcm2-7<sup>2C</sup>/Cdt1 expression strain | *yMH09 his::pST048(GAL1,10-MCM2-LPETGG, Flag-MCM3 ) lys2::pSKM002 (GAL1,10-MCM4,MCM5) trp::pSKM003 (GAL1,10-MCM6,MCM7) ura::pALS1(GAL1,10-Cdt1,GAL4)* | (Ticau et al., 2017) |
| yST180 | Mcm2-7<sup>4N</sup>/Cdt1 expression strain | *yMH09 his::pSKM004(GAL1,10-MCM2,Flag-MCM3) lys2::pST034(GAL1,10-UbSORT-MCM4,MCM5) trp1::pSKM003 (GAL1,10-MCM6,MCM7) ura::pALS1(GAL1,10-Cdt1,GAL4)* | This Study |
| ySG012 | Mcm2-7<sup>2C</sup> expression strain | *yMH09 his::pST048(GAL1,10-MCM2-LPETGG, Flag-MCM3 ) lys2::pSKM002 (GAL1,10-MCM4,MCM5) trp::pSKM003 (GAL1,10-MCM6,MCM7)* | This Study |
| ySG024 | Mcm2-7<sup>3N</sup> expression strain | *yMH09 his::pSG13 (GAL1,10-MCM2, Flag-TEV-GG-MCM3 ) lys2::pSKM002 (GAL1,10-MCM4,MCM5) trp::pSKM003 (GAL1,10-MCM6,MCM7)* | This Study |
| ySG032 | Mcm2-7<sup>2C,5RA</sup> expression strain | *yMH09 his::pST048(GAL1,10-MCM2-LPETGG, Flag-MCM3 ) lys2::pSKM028 (GAL1,10-MCM4,MCM5RA) trp::pSKM003 (GAL1,10-MCM6,MCM7)* | This Study |
| ySG033 | Mcm2-7<sup>3N,5RA</sup> expression strain | *yMH09 his::pSG13 (GAL1,10-MCM2, Flag-TEV-GG-MCM3 ) lys2::pSKM028(GAL1,10-MCM4,MCM5RA) trp::pSKM003 (GAL1,10-MCM6,MCM7)* | This Study |
| ySG10 | ORC<sup>5C</sup> expression strain | *yRH101 ura::pJF19 (Gal1,10- CBP-Orc1, Orc2); his3::pJF17 (Gal1,10-Orc3, Orc4); trp1::pSG01 (Gal1,10-Orc5-LPETGG, Orc6)* | This Study |
| yAZ55 | ORC<sup>1N</sup> expression strain | *yRH101 ura::pAA01 (UbsORT-FLAG-SDOrc1-Gal-SDOrc2); his3::pJF17 (SDOrc3-Gal-SDOrc4); trp1::pJF18 (SDOrc5-Gal-SDOrc6)* | This Study |
| ySG040 | ORC<sup>6C</sup> expression strain | *yRH101 ura::pJF19 (Gal1,10- CBP-Orc1, Orc2); his3::pJF17 (Gal1,10-Orc3, Orc4); trp1::pAZ63 (Gal1,10-Orc5, Orc6-LPETGG)* | This Study |
| ySG041 | ORC<sup>6C,6ΔN119</sup> expression strain | *yRH101 ura::pJF19 (Gal1,10- CBP-Orc1, Orc2); his3::pJF17 (Gal1,10-Orc3, Orc4); trp1::pSG21 (Gal1,10-Orc5, 3X-Flag-Δ119-Orc6-LPETGG)* | This Study |
| yST103 | Cdt1<sup>N</sup> expression strain | *yMH09 ura::pST013(GAL1,10-Ubsort-Cdt1-Flag, GAL4)* | This Study |
| ySG046 | Cdt1<sup>C</sup> expression strain | *yMH09 ura::pSG26(GAL1,10-3x-Flag-Cdt1-LPETGG, GAL4)* | This Study |

**Supplementary Table 2.**
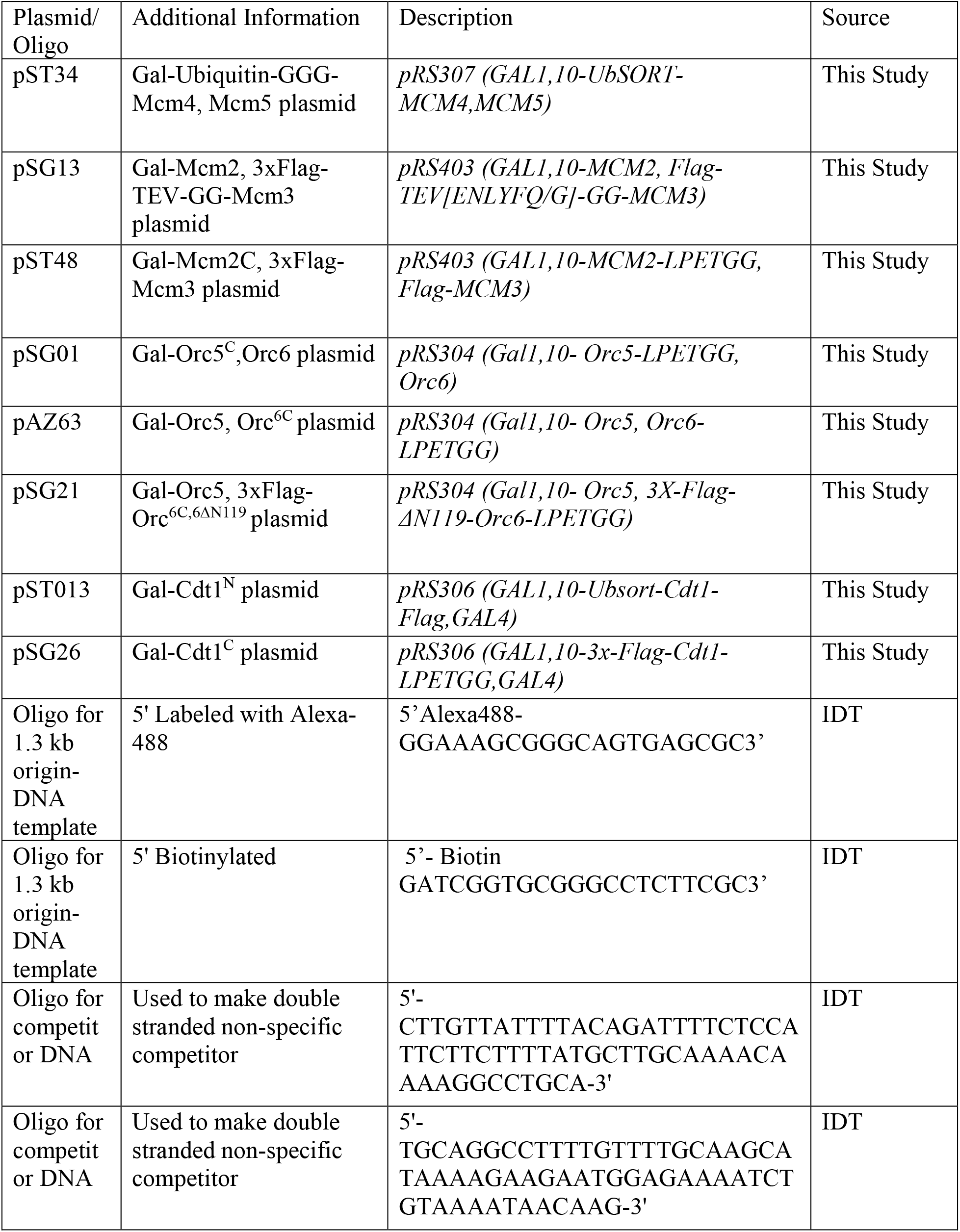
Plasmids and Oligos Used in this Study.

